# Distinct and additive effects of calorie restriction and rapamycin in aging skeletal muscle

**DOI:** 10.1101/2021.05.28.446097

**Authors:** Daniel J. Ham, Anastasiya Börsch, Kathrin Chojnowska, Shuo Lin, Aurel B. Leuchtmann, Alexander S. Ham, Marco Thürkauf, Julien Delezie, Regula Furrer, Dominik Burri, Michael Sinnreich, Christoph Handschin, Lionel A. Tintignac, Mihaela Zavolan, Nitish Mittal, Markus A. Rüegg

## Abstract

As global life expectancy continues to climb, maintaining skeletal muscle function is increasingly essential to ensure a good life quality for aging populations. Calorie restriction (CR) is the most potent and reproducible intervention to extend health and lifespan, but is largely unachievable in humans. Therefore, identification of “CR mimetics” has received much attention. CR targets nutrient-sensing pathways centering on mTORC1. The mTORC1 inhibitor, rapamycin, has been proposed as a potential CR mimetic and is proven to counteract age-related muscle loss. Therefore, we tested whether rapamycin acts via similar mechanisms as CR to slow muscle aging. Contrary to our expectation, long-term CR and rapamycin-treated geriatric mice display distinct skeletal muscle gene expression profiles despite both conferring benefits to aging skeletal muscle. Furthermore, CR improved muscle integrity in a mouse with nutrient-insensitive sustained muscle mTORC1 activity and rapamycin provided additive benefits to CR in aging mouse muscles. Therefore, RM and CR exert distinct, compounding effects in aging skeletal muscle, opening the possibility of parallel interventions to counteract muscle aging.

## Introduction

Dietary or calorie restriction (CR) is the most potent and reproducible intervention to extend lifespan. However, CR as a human lifestyle is considered largely unachievable based on the sheer willpower required to maintain a low-calorie diet along with potential side effects including extreme leanness and cold sensitivity (*1*). In order to eventually translate the health benefits of CR into medical treatments for aging humans, a mechanistic understanding of how CR confers longevity is essential (*2*). CR, along with many of the interventions known to prolong lifespan, dampen the activity of nutrient-sensing pathways, which center around the mammalian target of rapamycin complex 1 (mTORC1), thereby alleviating protein synthetic burden and promoting intrinsic quality control processes, like autophagy (*3*). Findings that the mTORC1 inhibitor, rapamycin (RM), extends lifespan in yeast (*4*), flies (*5*), worms (*6*) and mice (*7*) strengthened the hypothesis that mTORC1 inhibition is fundamental to CR-induced lifespan extension (*8*).

Over the last century, global life expectancy has nearly doubled, but, as the World Health Organization recognized while declaring 2021-2030 the decade of healthy aging: “adding more years to life can be a mixed blessing if it is not accompanied by adding more life to years” (*9*). A well-functioning neuromuscular system is fundamental to ensure more life comes with more years. Since anabolic pathways, especially those involving mTORC1, promote both muscle growth (*10, 11*) and sarcopenia, the age-related loss of muscle mass (*12*), skeletal muscle is considered a potential sticking point for achieving both life extension and life quality via CR and RM. However, we have recently demonstrated that long-term RM treatment is overwhelmingly, although not entirely, beneficial for aging mouse skeletal muscle (*13*). While a thorough examination of whether long-term CR, spanning the time of sarcopenic development (20-28 months in mice (*14*)) counteracts phenotypic and molecular signatures of sarcopenia is still lacking, life-long CR appears to afford similar benefits as RM to aging skeletal muscle, including a more stable neuromuscular junction (*15*) and slower age-related muscle loss (*16, 17*). Could RM, therefore, function as a CR mimetic to slow muscle aging?

Studies in mice highlight that although both CR and RM blunt mTORC1 activity and promote autophagy (*18, 19*), their effects on insulin signaling strongly diverge, with CR improving and RM impairing glucose tolerance (*19, 20*). Similarly, molecular profiling studies in mouse liver have shown that the vast majority of acute transcriptomic, metabolomic (*21*) and proteomic responses to CR and RM are distinct (*22*). But is this a case of ‘all roads lead to Rome’, where CR and RM travel different paths to mTORC1 suppression, or do these two quintessential life-prolonging interventions travel different roads with distinct destinations? If the former is true, we reasoned that 1) the molecular and phenotypic signatures of long-term CR and RM in skeletal muscle should overlap; 2) the beneficial effects of CR should be lost if skeletal muscle mTORC1 activity remains high and; 3) the effects of RM should be lost in calorie-restricted mice.

Using repeated measurements of whole-body muscle function and body composition, multi-muscle gene expression profiling and extensive endpoint examination of isolated muscle function and fiber type properties, we first thoroughly characterized the impact of long-term CR starting from 15 or 20 months until 30 months of age, thus covering the period when sarcopenia develops. Compared to a weight-controlled, *ad libitum*-fed control group, CR mice improved whole-body and isolated relative muscle function and experienced a fast-to-slow shift in muscle fiber phenotype. While much of this phenotype was shared by RM treated mice (*13*), gene expression signatures of CR and RM were strikingly divergent, the only exception relating to the suppression of age-related increases in immune and inflammatory responses. In further opposition to overlapping functions, CR improved markers of muscle quality, including autophagy blockade as seen by P62 build up, plasma creatine kinase (CK) levels, centro-nucleated fibers and *in vitro* muscle function in a nutrient-insensitive, mTORC1-driven model of premature muscle aging, without suppressing mTORC1 activity. Most conclusively, long-term RM treatment effectively counteracted skeletal muscle aging in both *ad libitum* and CR mice. We therefore demonstrate that RM and CR exert distinct and frequently additive effects on aging skeletal muscle, thereby opening the possibility of parallel interventions to counteract aging.

## RESULTS

### Adaptations to calorie restriction favor whole-body muscle function but do not prevent an age-related decline

After habituation to single housing and the AIN-93M diet, 15-(CR_15m_) or 20-(CR_20m_) month-old mice underwent a weekly, stepwise reduction in food intake from 100% (3.2g) to 90% (2.9g), 80% (2.6g), 70% (2.3g) and finally 65% (2.1g) of *ad libitum* levels until 30 months of age (Fig. 1A). After the initial month, calorie restricted mice continued to rapidly lose body mass for 2-3 months and then maintained a body mass between 21 and 26% below control mice from ∼19 or 23 months to 30 months for CR_15m_ and CR_20m_ groups, respectively (Fig. 1B). Food intake normalized to body surface area indicated that calorie restricted mice do not fully compensate for reduced food intake through body mass reductions, partially adapting to the low-energy environment through energy sparing processes (Fig. 1C). After initially drawing on energy reserves, particularly fat (*23*), calorie restricted mice display a range of adaptations to cope with the shortfall in energy intake, including improving energy absorption, reducing non-essential organ mass and lowering body temperature and energy expenditure (*24*). Compared to controls, 24-month-old calorie restricted mice had larger proportional differences in fat mass than lean mass, but smaller absolute differences (Fig. 1D). A progressive, age-related loss of whole-body fat mass in control mice narrowed the difference in fat, but not lean mass between CR and CON mice at 30 months of age.

**Figure 1:**
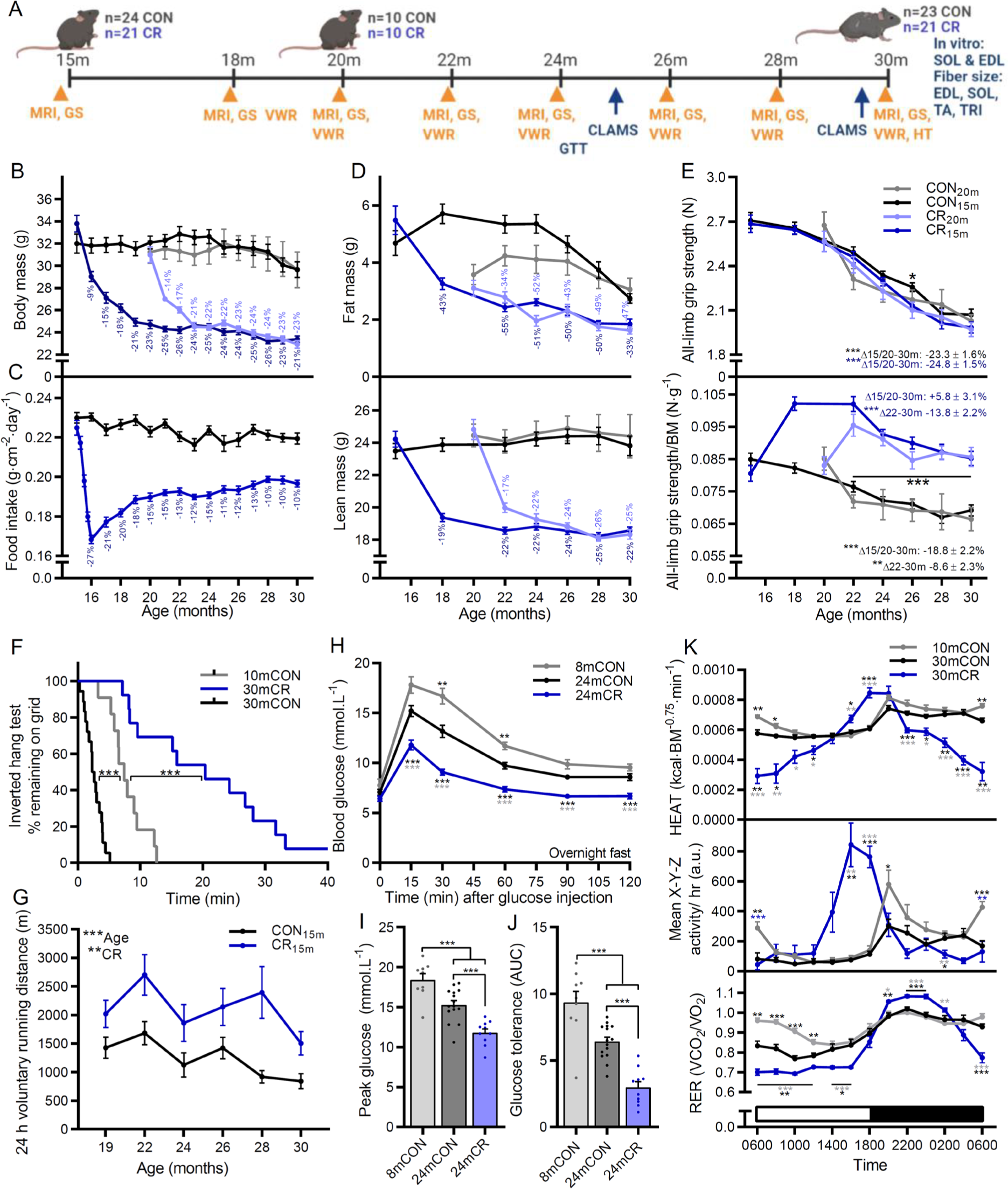
CR promotes beneficial metabolic and functional adaptations but does not prevent the age-related loss of muscle function. **(A)** Experimental design schematic showing start and endpoints of middle and late intervention groups as well as time course of physiological measures including body composition (MRI), grip strength (GS), voluntary wheel running (VWR), whole-body metabolism (CLAMS), hang test (HT) and glucose tolerance test (GTT). **(B)** Body mass for mouse groups fed *ad libitum* or 65% of *ad libitum* beginning at 15 months (CON_15m_ and CR_15m_) or 20 months (CON_20m_ and CR_20m_) of age. **(C)** Mean daily food intake normalized to body surface area for middle-aged groups. **(D)** Bimonthly recordings of whole-body fat (upper) and lean mass (lower); *n* = 18 (CON_15m_), 13 (CR_15m_), 6 (CON_20m_), and 8 (CR_20m_) mice. **(E)** absolute (upper) and body mass normalized (lower) all-limb grip strength; *n* = 19 (CON_15m_), 14 (CR_15m_), 6 (CON_20m_), and 8 (CR_20m_) mice. **(F)** Kaplan–Meier plot for the inverted grid-hang test performed prior to endpoint measures at 30 months of age for the middle-aged group; *n* = 11 (10mCON), 18 (CON_15m_), and 13 (CR_15m_) mice. **(G)** Twenty-four hours of voluntary running-wheel distance; *n* = 16 (CON_15m_), 13 (CR_15m_), 6 (CON_20m_), and 9 (CR_20m_) mice. Glucose tolerance test parameters including **(H)** blood glucose response to 2 mg·kg^-1^ glucose injection (I.P.), **(I)** peak glucose and **(J)** area under the curve/glucose tolerance. **(K)** Whole-body metabolic analysis of energy expenditure normalized to body surface area (upper), mean X-Y-Z activity (middle) and respiratory exchange ratio (VCO_2_/VO_2_; lower) reported every 2 h across one full day (white)/night (black) cycle in the month prior to endpoint measures; *n* = 12 (10mCON), 9 (30mCON), and 7 (30mCR) mice. Data are presented as mean ± SEM. Two-way repeated-measure ANOVA with Tukey post hoc tests (B–E, G-H, K), Mantel–Cox log rank (F), and one-way ANOVA with Fisher’s LSD post hoc tests (I-J) was used to compare the data. *, **, and *** denote a significant difference between groups of *P* < 0.05, *P* < 0.01, and *P* < 0.001, respectively. Colored asterisks refer to the group of comparison.

Repeated all-limb grip strength measures spanning the treatment period show that the progressive age-related loss of absolute grip strength was similar in both control and calorie restricted mice (P<0.001; Fig. 1E, upper). However, due to the initial rapid loss of body mass, calorie restricted mice strongly increased relative grip strength (N·g body mass^-1^) over the initial 2-3 months before displaying a similar age-related decline in relative grip strength as control mice from 22 to 30 months (Fig. 1E, lower). As such, compared to the initial recording (15 or 20 months), relative grip strength significantly declined across the treatment period in control mice (P<0.001), but not in CR mice. However, when compared to the 22-month recording, the decline in relative grip strength was not different between control (P<0.002) and calorie restricted (P<0.001) mice at 30 months. Consistent with improved relative strength, calorie restriction potently improved inverted grid-hang time compared to both 10mCON and 30mCON groups (Fig. 1F; P<0.001). Similarly, voluntary running activity over a 24 h period was consistently improved in calorie restricted mice across the trial (Fig. 1G; P<0.01).

After an overnight fast for all groups, calorie restricted mice showed a marked increase in glucose tolerance and significantly lower blood glucose levels than 8 and 24-month-old control mice at each point after glucose injection (Fig. 1H). CR also lowered peak glucose (Fig. 1I) and the area under the curve (AUC; Fig. 1J). Despite similar body mass and whole-body fat mass, 24-month-old control mice showed lower blood glucose at 30 and 60 min after glucose injection as well as a lower peak glucose and AUC than 8-month-old control mice. Similarly, improvements in glucose tolerance have been observed between 20 and 28 months of age in wild type mice (*25*) and also in mice genetically modified to have high muscle fiber mTORC1 activity (*26*), a commonly observed feature of sarcopenic muscle (*13, 27–29*).

While daylight regulated whole-body metabolism in control mice; with energy expenditure, activity and RER all high at night and low during the day; metabolism in CR mice centered around food availability (Fig. 1K and Fig. S1A-F). In CR mice, energy expenditure (Fig. 1K upper) and activity (Fig. 1K middle) were tightly restricted to the hours immediately before and immediately after their 5 pm food allocation. Despite displaying characteristic anticipatory increases in activity and energy expenditure prior to food, RER remained at around 0.7 (Fig. 1K lower), indicating almost exclusive fat utilization, before a food- and therefore glucose availability-related increase during night-time hours. In line with lower food intake per body surface area in CR mice (Fig. 1C), mean energy expenditure per body surface area was - 8.8 and -8.5% lower (P<0.05) during day-time hours and -16.6% and -21.0% lower during night-time hours at 25 and 30 months of age in CR than control mice (Fig. S1C). Together, these data indicate that calorie restricted mice shed excess tissue mass and induce metabolic adaptations to minimize energy use and maximize energy uptake when scarce food becomes available. Since the extent of metabolic adaptations and lifespan extension increase in parallel with that of calorie restriction, these adaptations are thought to be central to the health and longevity benefits of CR (*24, 30*). While these changes initially lead to a substantial increase in relative grip strength, calorie restricted mice also display a progressive age-related loss of grip strength indicating that while CR clearly improves muscle functional parameters, mice are not entirely spared from muscle aging.

### Calorie restriction promotes a fast-to-slow muscle phenotype transition

Aging reduced absolute (Fig. S1G) and relative (mg/g body mass) mass (Fig. 2A) in all measured muscles. CR further reduced absolute muscle mass in the fast-twitch quadriceps (QUAD), gastrocnemius (GAS), tibialis anterior (TA), plantaris (PLA), extensor digitorum longus (EDL) and triceps brachii (TRI) muscles but not in the slow-twitch SOL muscle (Fig. S1G). When accounting for body mass, muscle mass was similar in 30mCON and 30mCR groups for all fast-twitch muscles, except quadriceps, which along with the slow-twitch soleus muscle were significantly heavier relative to body mass in 30mCR than 30mCON mice (Fig. 2A). While correlation graphs indicate that the muscle-to-body mass ratio was similar for all fast twitch muscles, SOL mass was clearly higher for a similar body mass in 30mCR than 30mCON mice (Fig. 2B). In line with measures of body composition, CR reduced the absolute mass of all major organs (heart, spleen, kidney, liver) as well as epididymal white and interscapular brown fat depositions and prevented the age-related accumulation of non-functional mass (i.e. seminal vesicles; Fig. S1H). Proportional to body mass, CR preserved heart, liver and brown fat mass while shedding white fat, kidney and spleen mass (Fig. S1I).

**Figure 2:**
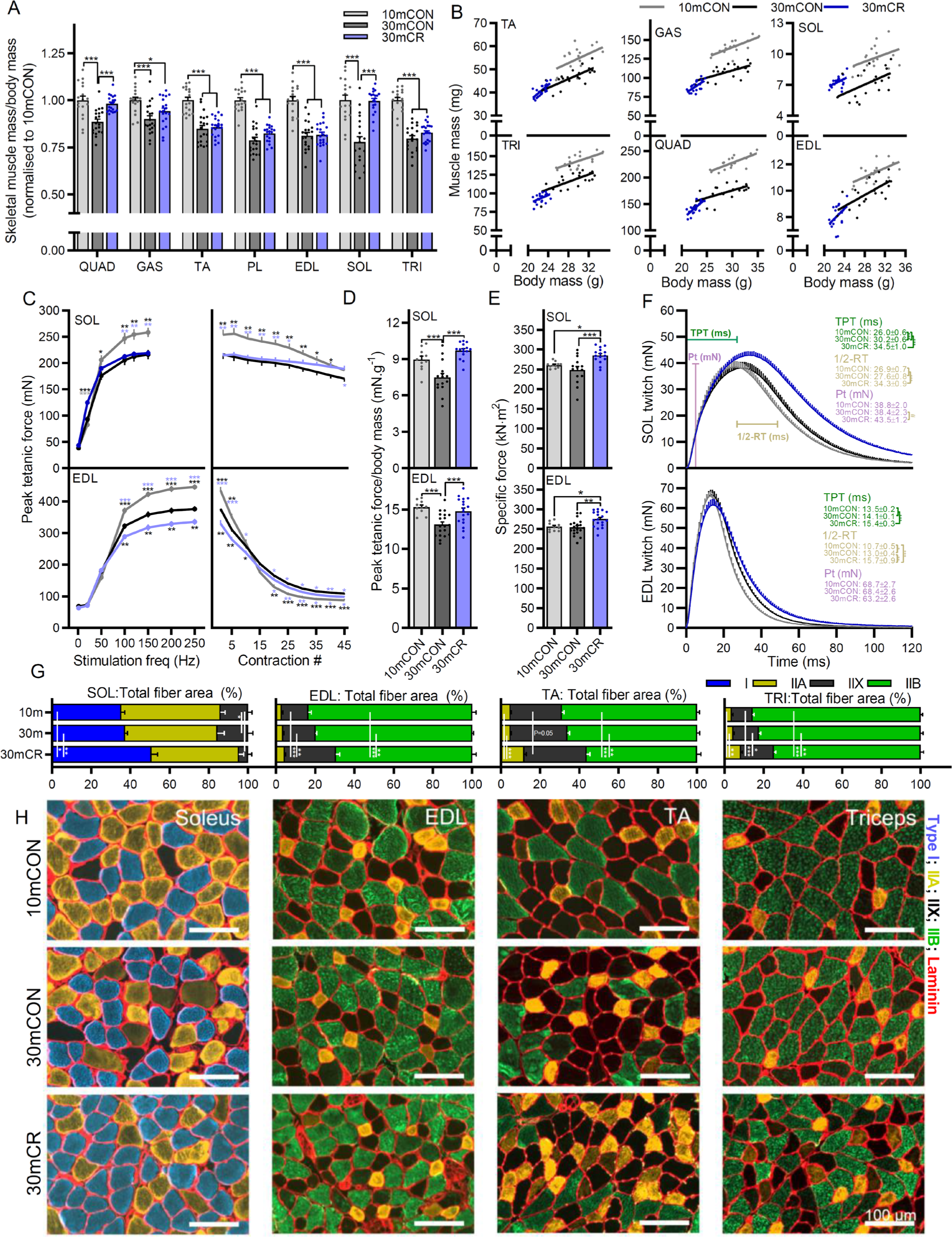
CR promotes a fast-to-slow muscle fiber phenotype shift. **(A)** Muscle mass for *quadriceps* (QUAD), *gastrocnemius* (GAS), *tibialis anterior* (TA), *plantaris* (PLA), *extensor digitorum longus* (EDL), *soleus* (SOL), and *triceps brachii* (TRI) was averaged across both limbs, normalized to body mass and then to 10-month-old control mice. **(B)** Scatterplots and linear regressions of the relationship between body and muscle mass of the fast twitch TA, TRI, GAS, QUAD and EDL muscles and the slow twitch SOL muscle. Isolated muscle function parameters, including **(C)** force-frequency curve (left) and fatigue response to multiple stimulations (right), **(D)** peak force normalized to body mass, **(E)** peak force normalized to cross sectional area (specific force), and **(F)** mean twitch responses including time-to-peak tension (TPT), half-relaxation time (1/2-RT) and peak twitch (Pt) for SOL (top panel) and EDL muscle (bottom panel). **(G)** Proportional total fiber-type-specific cross-sectional area analyzed on whole cross sections of (left to right) SOL (n=6), EDL (n=7, 9 and 8), TA (n=11, 13 and 7) and TRI (n=5, 9 and 9) for 10mCON, 30mCON and 30mCR, stained with antibodies against type I (blue), type IIA (yellow), and type IIB (green) fibers as well as laminin (red), while fibers without staining were classified as IIX. **(H)** Representative images for SOL, EDL, TA and TRI. Group numbers for 10mCON are *n* = 17 (a, b), 10 (c-f: EDL) and 8 for fatigue, 11 (c-e: SOL) or 9 for fatigue, for 30mCON *n* = 20 (a, b), 19 (c–f: EDL), 15 (c–e: SOL) and 16 (f: SOL), and for 30mCR *n* = 20 (a, b), 18 (c–e: EDL), 15 (c-e: SOL) or 13 for fatigue, 12 (f: SOL) and 15 (f: EDL). Data are presented as mean ± SEM. One-way (a and d-f) or two-way repeated-measure (c, g) ANOVAs with Fisher’s LSD or Tukey’s post hoc tests, respectively, were used to compare between data. *, **, and *** denote a significant difference between groups of *P* < 0.05, *P* < 0.01, and *P* < 0.001, respectively. Colored asterisks refer to the group of comparison.

Preferential sparing of slow-twitch muscle was also observed in measures of isolated muscle function in the fast EDL and slow SOL muscle (Fig. 2C-F). Long-term CR did not alter the strong age-related reduction in absolute tetanic force in the SOL but further lowered tetanic force in EDL muscle from 100-250 Hz (Fig. 2C, left). In response to repeated stimulations, tetanic force dropped more rapidly in the 10mCON group than either 30mCON or 30mCR groups (Fig. 2C, right). Despite producing significantly greater absolute force at the start of the assay in both SOL and EDL, force in the 10mCON was not different to the other groups after 45 contractions in SOL and significantly lower than both other groups after 20 contractions in the EDL (Fig. 2C, right). After 45 contractions, 30mCR mice produced significantly more force than 30mCON mice in SOL muscle, but less in EDL muscle. In line with body mass-normalized grip-strength measurements, calorie restriction completely prevented the age-related loss of SOL and EDL peak tetanic force (Fig. 2D). This was at least partially the result of improved muscle quality in 30mCR mice, which showed significantly higher specific muscle force, representing peak force normalized to muscle cross-sectional area, in both SOL and EDL muscle (Fig. 2E). Aging slowed muscle twitch properties with significantly longer time-to-peak tension in SOL and a longer half-relaxation time in the EDL muscle (Fig. 2F). CR augmented this phenotype, with 30mCR mice showing significantly longer time-to-peak tension and half-relaxation time than either 10mCON or 30mCON in both SOL and EDL muscles (Fig. 2F).

Consistent with the CR-induced fast-to-slow shift in muscle function properties, preferential protection of slow-twitch muscle mass, and improvements in whole-body endurance, fiber type-specific analysis of muscle cross sections showed a strong increase in the proportion of total cross-sectional area made up by slower fiber types in soleus (SOL), extensor digitorum longus (EDL), tibialis anterior (TA) and triceps brachii (TRI; Fig. 1G-H). Although a fast-to-slow transition was seen in all muscles analyzed, each muscle displayed specific changes in fiber type size and number (Fig. S2A-E). Compared to 10mCON and 30mCON, the proportional area of type I fibers in SOL, IIX fibers in EDL, IIA fibers in TA and both IIA and IIX fibers in TRI was higher while the proportional area of IIX in SOL and IIB in EDL, TA and TRI was lower for 30mCR (Fig. 1G-H). The largest and fastest type IIB fibers appeared to be the most expendable for calorie restricted mice, with significant reductions in both proportional fiber number (Fig. S2B) and minimum fiber feret (Fig. S2C-D) in EDL, TA and TRI muscles. On the other hand, the size of slower IIA fibers and of intermediate IIX fibers were less affected by both age and CR (Fig. S2C-D) and CR increased their proportional number (Fig. S2B). Since slower-type fibers are more resistant to age-related atrophy, the CR-induced fast-to-slow fiber-type transition, despite the absolute decrease in muscle mass, may ultimately preserve muscle function in aging mice as seen with other interventions (*31*).

### Calorie restriction and rapamycin induce distinct transcriptomic changes

While the overt aging phenotype in calorie-restricted mice is distinct from that observed in mice treated with rapamycin over the same period (*13*), calorie restriction is thought to counteract aging in large part by suppressing nutrient-induced activation of mTORC1 activity (*32*). If this were the case, then long-term CR and RM should induce a common core gene expression signature in aging muscle. To address this hypothesis, we performed mRNA-sequencing on four muscles (GAS, TA, TRI and SOL) from six 30-month-old calorie restricted mice and compared the profiles with our previously reported 10mCON, 30mCON and 30mRM data sets (Fig. 3A) (*13*).

**Figure 3:**
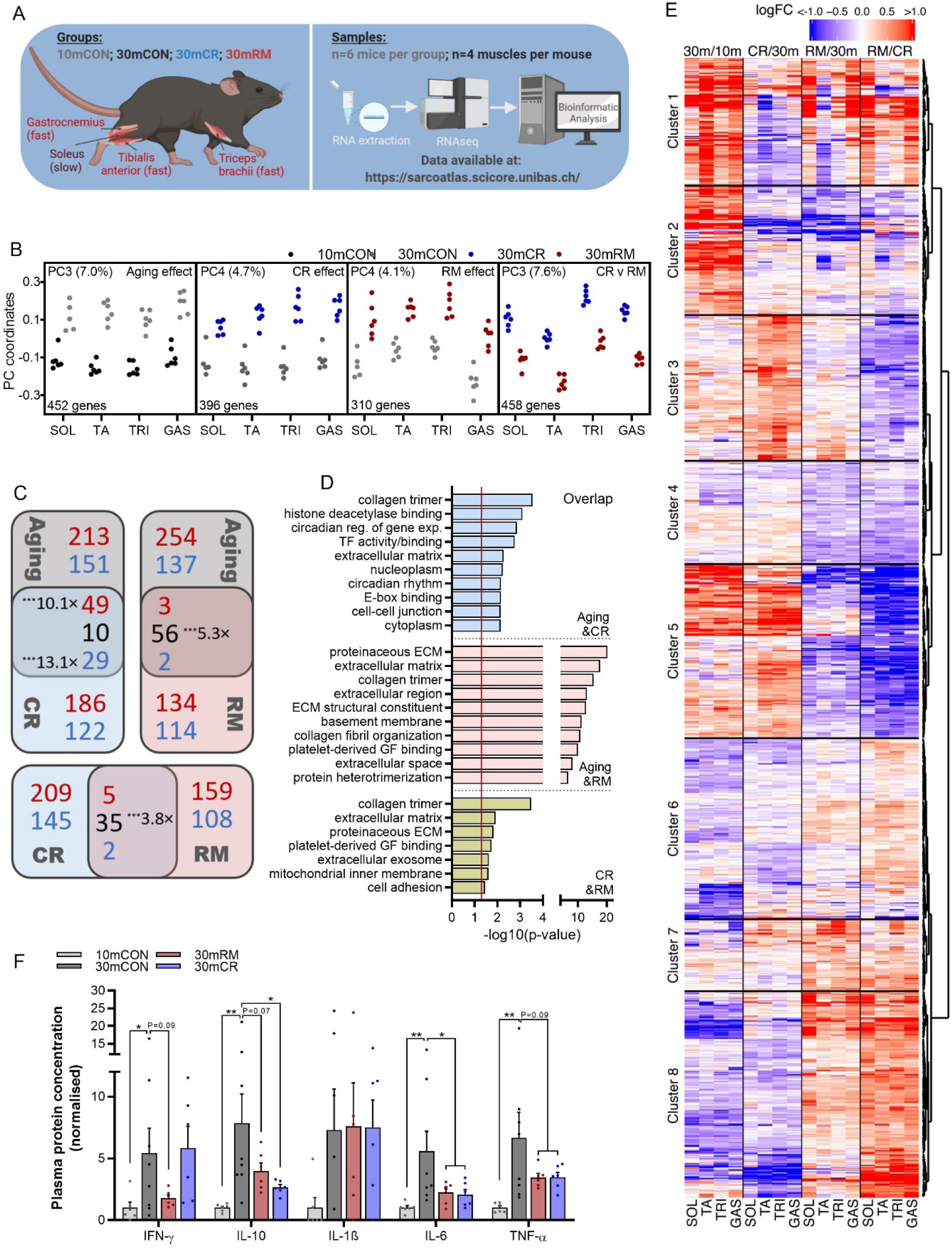
CR and RM induce distinct gene expression signatures. **(A)** Scheme of treatment groups, muscles, and numbers of samples used in sequencing analysis. Data for 10mCON, 30mCON, and 30mRM have been previously reported (Ham & Börsch et al. 2020). **(B)** Coordinates of principal components representing aging (PC3 for 10mCON and 30mCON) CR (PC4 for 30mCON and 30mCR), RM (PC4 for 30mCON and 30mRM) and CR vs. RM (PC3 for 30mCR and 30mRM) effects for gene expression collected in *soleus* (SOL), *tibialis anterior* (TA), *triceps brachii* (TRI) and *gastrocnemius* (GAS). The numbers associated with the PCs indicate the fraction of the variance in gene expression in samples along the corresponding PC. Each dot corresponds to one muscle sample, from an individual animal. The number of genes aligned with each PC is displayed in the bottom left corner of each graph. A gene was considered aligned with a PC if the absolute value of the Pearson correlation coefficient between the expression of the gene and PC coordinates was ≥0.4, and the absolute value of the *z* score of the projection of the gene expression on a PC was ≥1.96. **(C)** Pairwise Venn diagram comparisons of genes significantly aligned to aging, CR and RM effects. Numbers in red, blue and black represent increasing, decreasing and oppositely regulated genes, respectively. Where the overlap of genes is significantly above that expected by chance, the level of significance and representation factor are noted. **(D)** Top-ten DAVID gene ontology terms enriched (*P* < 0.05) for genes aligned to both aging and CR effects, aging and RM effects or CR and RM effects. Enrichment significance threshold was set at *P* < 0.05 (gray and red dashed lines). **(E)** Heatmap of fold-changes for genes aligned with any of the four PCs described in (B) for aging (30mCON/10mCON), CR (30mCR/30mCON), RM (30mRM/30mCON) and CR vs. RM (30mRM/30mCR) effects in all four muscles. Hierarchical clustering based on the Euclidean distance of these changes rendered 8 gene clusters. **(F)** Plasma cytokine protein concentration. Data are displayed as fold-change from 10mCON group. Cytokine levels between the detection limit were set as 0. For the sarcopenia data set, *n* = 6 mice per muscle per group, except for SOL 30mCON where one data point was removed due to a technical error. A modified Fisher’s exact test was used to determine significance.

To determine the overlap in core gene expression signatures induced by age, CR and RM, we first defined the specific effect of each condition separately by performing principal component (PC) analysis on data from all four muscles for each of the following four comparisons: 10mCON vs. 30mCON (PC_10m-30m_); 30m vs. 30mCR (PC_30m-CR_); 30m vs. 30mRM (PC_30m-RM_;) and; 30mCR vs. 30mRM (PC_CR-RM_;). Aside from three prominent PC patterns representing the inter-muscle gene expression differences we have previously reported (*13*), we found that for each comparison there was one PC (PC3 or PC4) that showed a clear between-group expression pattern difference common to all four muscles (Fig. 3B). To check to what extent genes aligned with PC_310m-30m_ (aging effect), PC_430m-CR_ (CR effect) and PC4_30m-RM_ (RM effect) overlapped, we created Venn diagrams for aging and CR, aging and RM and CR and RM effect comparisons, respectively (Fig. 3C). Strikingly, RM reversed and CR accentuated the vast majority of commonly regulated, age-related gene expression changes. Likewise, genes that responded to both CR and RM treatments were predominately regulated in the opposite direction. Gene ontology analysis showed that genes involved in extracellular matrix remodeling were highly represented in overlapping genes from each of the three comparisons (Fig. 3D).

Next we visualized the log-fold changes in expression for each gene aligned to any of the PCs displayed in Fig. 3B. Hierarchical clustering based on the Euclidean distance of these changes rendered 8 gene clusters with distinct gene expression patterns (Fig. 3E). While CR and RM both tended to suppress a group of genes related to the innate immune response which are upregulated with age in cluster 2, CR and RM displayed strikingly opposing gene expression patterns throughout most other clusters as well as distinct CR (cluster 3) and RM (cluster 4) effects related to lipid metabolism and insulin signaling, respectively. While RM reversed, or partially reversed, age-related gene expression changes in many clusters, CR augmented age-related signaling such that gene expression differences between CR and RM were exaggerated. This pattern was particularly prevalent in cluster 5 and 8. Cluster 5 represents genes increased with age, further increased by CR and suppressed by RM, while cluster 8 displays the opposite pattern. The top DAVID gene ontology terms associated with cluster 5 relate to changes in metabolism while those associated with cluster 8 relate to the extracellular matrix (Fig. S3). The common effects of CR and RM observed in cluster 2 primarily relate to immune responses.

In line with these findings, circulating levels of the pro-inflammatory cytokines tended to be lower in both calorie-restricted and rapamycin-treated mice, with the exception of IFN-γ which tended to be suppressed by RM, but not CR (Fig. 1F).

These striking differences in gene expression patterns led us to wonder whether the signaling responses to CR and RM represent distinct paths to a common destination centered around immune and inflammatory suppression, i.e. an ‘all roads lead to Rome’ scenario, or whether CR and RM act on distinct muscle aging processes. If the latter is true, we reasoned that calorie restriction should benefit aging skeletal muscle independent of skeletal muscle mTORC1 suppression.

### Calorie restriction attenuates muscle mTORC1-driven premature sarcopenia without suppressing mTORC1

To address this hypothesis, we calorie restricted muscle-specific TSC1-knockout (TSCmKO) mice, which display sustained, nutrient-insensitive mTORC1 activation (*33*) and an accelerated aging phenotype (*13, 26*). At 3 months of age, TSCmKO mice display high muscle mTORC1 activity and impaired autophagy, but mild phenotypic features. From ∼6 to 12 months of age, TSCmKO display a progressive sarcopenia-like phenotype including muscle atrophy and weakness as well as accumulation of P62-labelled proteins and aggregates (*33*), which are also, to some degree, features of sarcopenia (*13, 34*). We therefore tested whether 7 months of calorie restriction starting at 3 months of age can ameliorate the muscle fiber mTORC1-driven sarcopenic phenotype in TSCmKO mice (Fig. 4A). CR induced similar changes in body (Fig. 4B), fat (Fig. 4C) and lean (Fig. 4D) mass in both TSCmKO and WT control mice and comparable with those observed in naturally aging mice (Fig. 1B, D). *Ad libitum*-fed TSCmKO mice (TSC-AL) were significantly lighter than their *ad libitum*-fed controls (WT-AL) at 9.5 months. This was primarily due to a divergence in fat depositions from 4.5 months, with WT-AL gaining (+18%; P<0.05) and TSC-AL losing (-25%; P<0.05) fat mass over the period (Fig. 4B). Body mass and composition were comparable in calorie restricted TSCmKO (TSC-CR) and WT (WT-CR) mice, aside from a tendency for lower fat mass in TSC-CR than WT-CR at 6.5 months (Fig. 4B). CR also induced the characteristic metabolic adaptations in patterns of energy expenditure (Fig. 4E) and RER (Fig. 4F) equally in both WT and TSCmKO mice and comparable with those observed in naturally aging mice (Fig. 1K).

**Figure 4:**
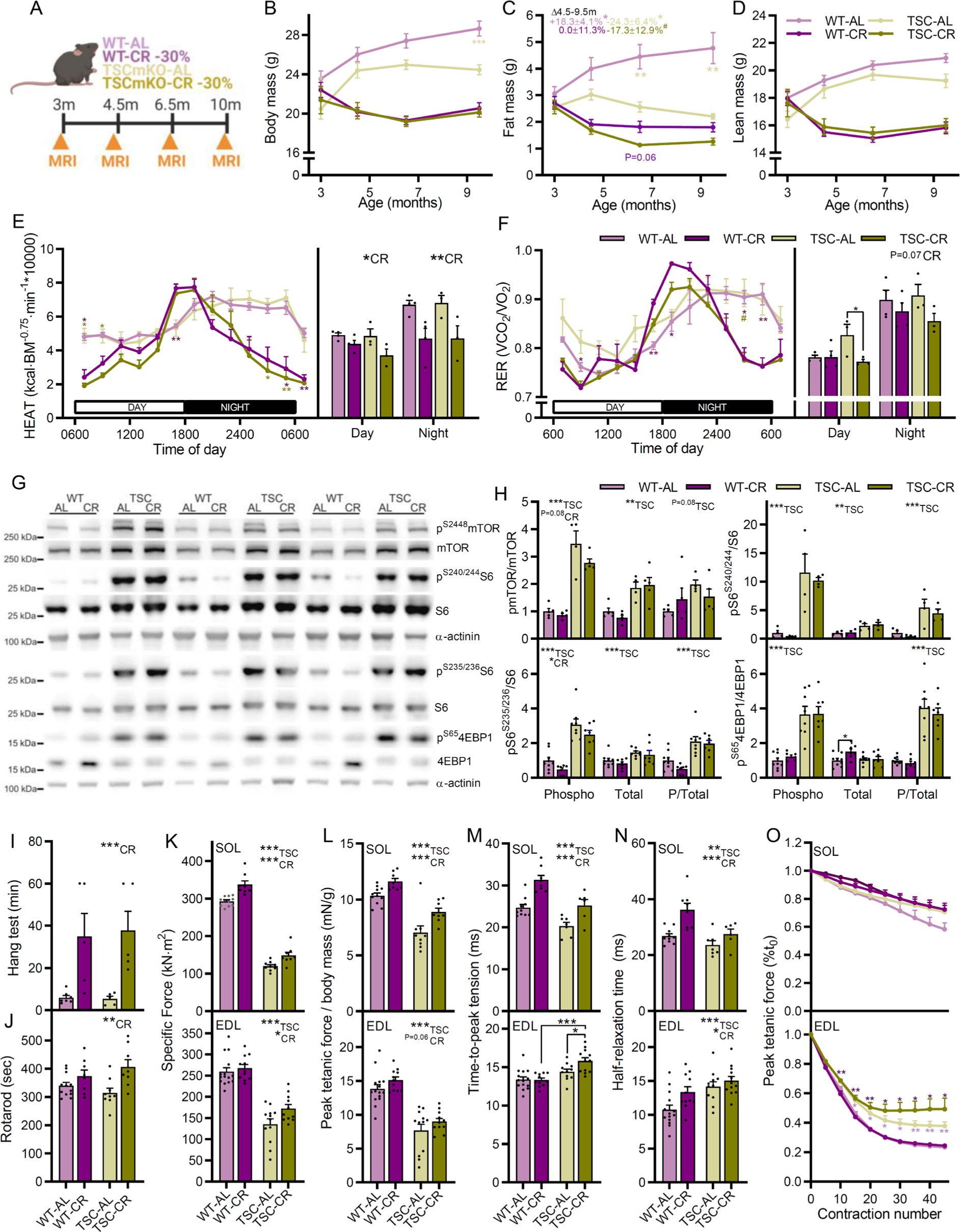
CR improves muscle function without suppressing mTORC1 activity in the TSCmKO model of accelerated muscle aging. **(A)** Experimental design schematic showing experimental groups as well as time course of physiological measures. **(B)** Body mass **(C)** whole-body fat mass and **(D)** lean mass for WT and TSCmKO mice fed *ad libitum* (WT-AL and TSC-AL) or 70% of *ad libitum* (WT-CR and TSC-CR) beginning at 3 months of age. **(E)** Whole-body metabolic analysis of energy expenditure normalized to body surface area and **(F)** respiratory exchange ratio (VCO_2_/VO_2_; lower) reported every 2 h across one full day (white)/night (black) cycle (left) and day and night-time averages (right) in the month prior to endpoint measures; *n* = 12 (10mCON), 9 (30mCON), and 10 (30mCR) mice. **(G)** Representative western blot analysis of mTORC1 pathway components in WT-AL, WT-CR, TSC-AL and TSC-CR *gastrocnemius* (GAS) muscle. Similar results were obtained for each protein across three separate gels with different samples. **(H)** Quantification of western blots showing the abundance of phosphorylated protein normalized to total protein for mTOR (upper left) S6 (upper right and lower left) and 4EBP1 (lower right). **(I)** Inverted grid hang time and **(J)** time spent on a rotating rod. Isolated muscle function parameters for SOL (upper panel) and EDL (lower panel), including **(K)** specific force, **(L)** peak tetanic force normalized to body mass, **(M)** twitch time-to-peak tension and **(N)** half-relaxation time as well as **(O)** fatigue response to multiple stimulations. Data are presented as mean ± SEM. Two-way ANOVAs with Tukey post hoc tests were used to compare the data. *, **, and *** denote a significant difference between groups of *P* < 0.05, *P* < 0.01, and *P* < 0.001, respectively. # denotes a trend where 0.05 < *P* < 0.10. Colored asterisks refer to the group of comparison.

To confirm that mTORC1 activity was not suppressed in TSC-CR mice, we investigated the phosphorylation status of mTOR (S2448) and its key downstream targets S6 (S240/244 and S235/237) and 4EBP1 (S65) as well as AKT (S473) and PRAS40 (T246), which are dampened by inhibitory feedback from the mTORC1 target S6K1 via IRS1 (*35*) in TA and/or GAS muscle using Western blot analysis (Fig. 4G-H and S4A-B & D). TSC-AL mice displayed the stereotypical markers of chronic mTORC1 activation, including high phosphorylated mTOR, S6 and 4EBP1 as well as lower phosphorylated AKT and PRAS40. Despite reducing the availability of mTORC1-activating nutrients (e.g. amino acids), CR was largely incapable of suppressing mTORC1 activity or alleviating feedback inhibition of AKT in TSC-CR mice (Fig. 4G-H and S4A-B, E). Together, these results indicate that CR induces typical changes in whole-body composition and metabolism independent of muscle mTORC1 activity and without overt negative side-effects.

Despite a lack of effect on mTORC1 activity, CR robustly improved whole-body measures of muscular endurance (Hang test, Fig. 4I) and coordination (rotarod, Fig. 4J), independent of genotype. Likewise, CR improved or tended to improve both specific force (Fig. 4K) and peak tetanic force normalized to body mass (Fig. 4L) in isolated SOL and EDL muscle in both WT and TSCmKO mice. As we observed in aging WT mice, CR induced a slowing of muscle twitch properties, particularly in the SOL muscle (Fig. 4M-N), although fatigability was not significantly affected (Fig. 4O).

As a result of chronic mTORC1 activity-induced phosphorylation of ULK1 and thereby autophagy blockade, old (9-12 months) TSCmKO mice display strong cytosolic and aggregated P62 accumulation in muscle fibers and an associated increase in signs of muscle degeneration, including high levels of plasma CK and centro-nucleated, regenerating fibers (*33*). Indeed, TSC-AL mice displayed a strong increase in P62 aggregate- and cytosolic-positive fibers (Fig. 5A-C), P62 protein accumulation in muscle lysates (Fig. 5D), plasma CK activity (Fig. 5E) and centro-nucleated fibers (Fig. 5F). Strikingly, CR almost completely alleviated P62 accumulation and signs of muscle degeneration. Despite the strong reduction in P62 protein levels, CR did not suppress *Sqstm1* expression, which encodes for the P62 protein, nor the other autophagy induction mediators *Ctsl*, *Gabarapl2*, *Bnip3*, *Atg7* or *Map1lc3a* (Fig. 5C). Likewise, the mTORC1-mediated phosphorylation of ULK1 (S757), the LC3II/LC3I ratio and protein levels of beclin1 and BNIP3 were all unaltered by CR (Fig. 5I-K).

**Figure 5:**
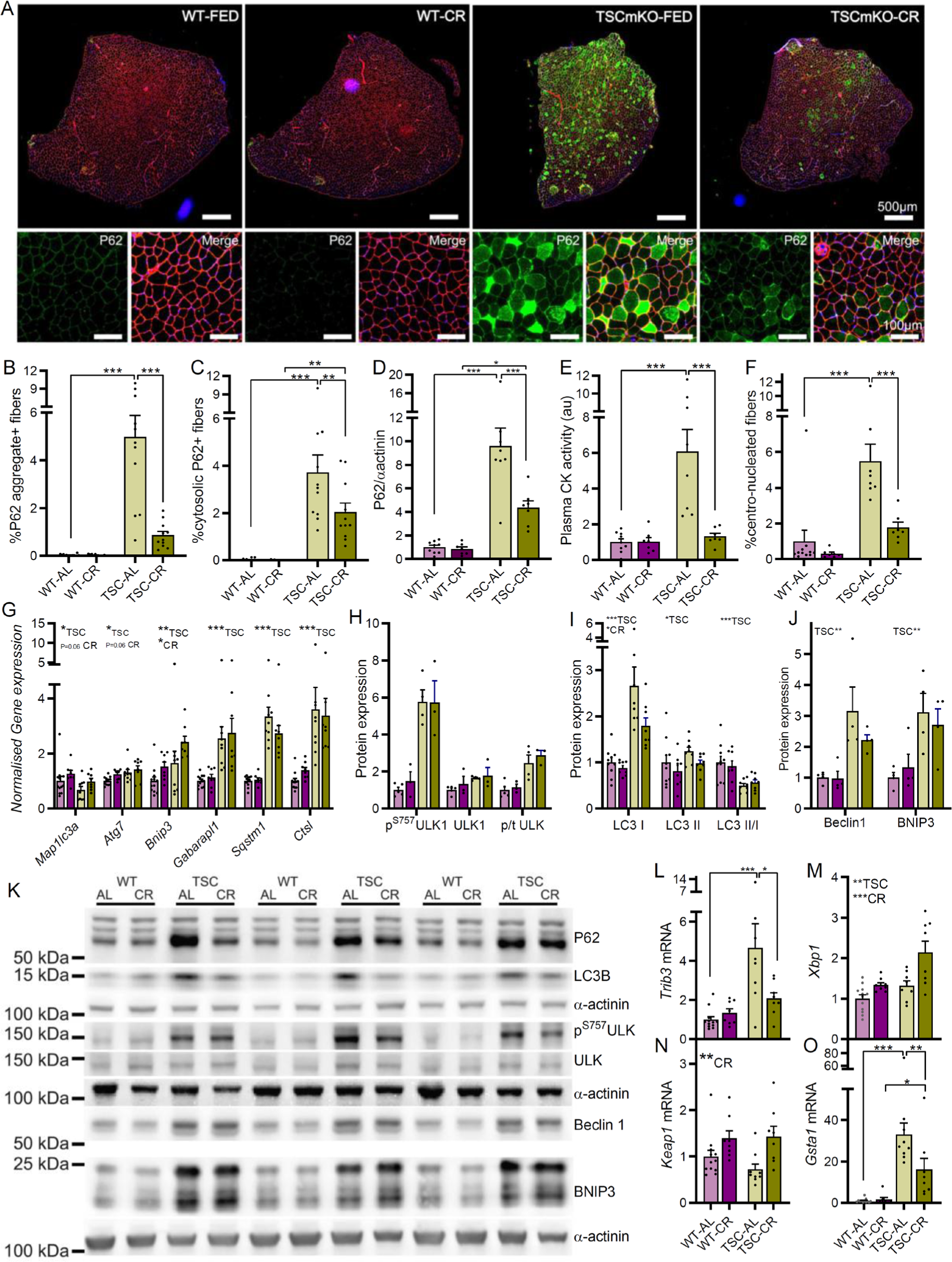
CR attenuates P62 accumulation and improves muscle integrity in TSCmKO mice. **(A)** Representative tibialis anterior (TA) cross sections stained with antibodies against P62 and laminin and counterstained with DAPI. Quantification of fibers with **(B)** P62+ aggregates and **(C)** P62+ cytosolic staining. **(D)** Western blot quantification of P62 protein expression in gastrocnemius muscle. **(E)** Plasma creatine kinase activity and **(F)** percentage centro-nucleated fibers in WT-AL, WT-CR, TSC-AL and TSC-CR mice. **(G)** RT-qPCR analysis of autophagy associated genes in gastrocnemius muscle. Western blot quantification of the abundance of **(H)** phosphorylated and total ULK1 protein, **(I)** LC3I, LC3II and the ratio of LC3II to I and **(J)** beclin1 and BNIP3 protein as well as **(K)** representative gels. RT-qPCR analysis of ER-stress and autophagy interacting genes including **(L)** Trib3, **(M)** Xbp1, **(N)** Keap1 and **(O)** Gsta1. Data are presented as mean ± SEM. Two-way ANOVAs with Tukey post hoc tests were used to compare data. *, **, and *** denote a significant difference between groups of *P* < 0.05, *P* < 0.01, and *P* < 0.001, respectively. # denotes a trend where 0.05 < *P* < 0.10. Colored asterisks refer to the group of comparison.

Endoplasmic reticulum (ER) stress and activation of the unfolded protein response (UPR) is also a hallmark of chronic muscle mTORC1 activation (*26*). CR did not reduce signs of ER stress in TSCmKO mice, including increased levels of ATF4, FGF21 and BiP (Fig. S4C-D). However, CR altered the response to ER stress, blunting the mTORC1-driven upregulation of *Trib3* (Fig. 5L), which can bind and inhibit the breakdown of P62, while inducing *Xbp1* (Fig. 5M), encoding a UPR-inducing transcription factor (*36*). CR also induced the expression of *Keap1* (Fig. 5N), a gatekeeper protein for the NRF2-induced stress response that can be sequestered by P62, thereby preventing it from binding and preventing NRF2 nuclear translocation. While initially cytoprotective, chronic NRF2-induced activation of the stress response is deleterious (*37*). In line with a CR-induced suppression of NRF2 activity by *Keap1* upregulation, the mTORC1-induced upregulation of *Gsta1* expression, an NRF2 target, was strongly suppressed in TSC-CR mice (Fig. 5O).

Together, these data strongly support the idea that CR can improve muscle proteostasis, integrity and function without suppressing mTORC1 activity. But is the reciprocal also true, or are the beneficial effects of rapamycin-induced mTORC1 suppression on aging skeletal muscle already captured by CR? In other words, would rapamycin also slow muscle aging in CR mice?

### CR and RM have distinct and often additive beneficial effects on aging skeletal muscle

To directly determine whether RM and CR exert non-overlapping effects in aging skeletal muscle, *ad libitum*-fed (CON) and 35% calorie restricted (CR) male C57BL/6 mice were fed a standardized AIN-93M diet containing either encapsulated rapamycin (RM and CR+RM) or an equivalent amount of encapsulating vehicle (Eudragit; CON and CR) starting from 19 months of age (Fig. 6A). Based on body mass and food intake data from our initial study and a target dose of ∼4 mg·kg^-1^·day^-1^ rapamycin, 42 and 48 mg·kg^-1^ of active encapsulated rapamycin was incorporated into the diet of RM and CR+RM groups, respectively. Consistent with our previous results, body mass loss compared to pre-trial levels was observed from 25 months in CON mice and from 20 months in RM-treated mice. After the initial adjustment to reduced food intake by 20 months of age, CR and CR+RM mice also displayed an age-related decline in body mass from around 26 months (Fig. 6B). Food intake, normalized to body surface area, was relatively stable over the treatment period in CON and RM mice, while CR and CR+RM mice showed a steady increase in normalized food intake from 20 months of age following an initial steep decline (Fig. 6C). RM did not alter the CR or age-related decline in whole-body fat mass and despite reducing lean mass in *ad libitum* fed mice, RM did not alter the CR-induced decline in whole-body lean mass (Fig. 6D).

**Figure 6.**
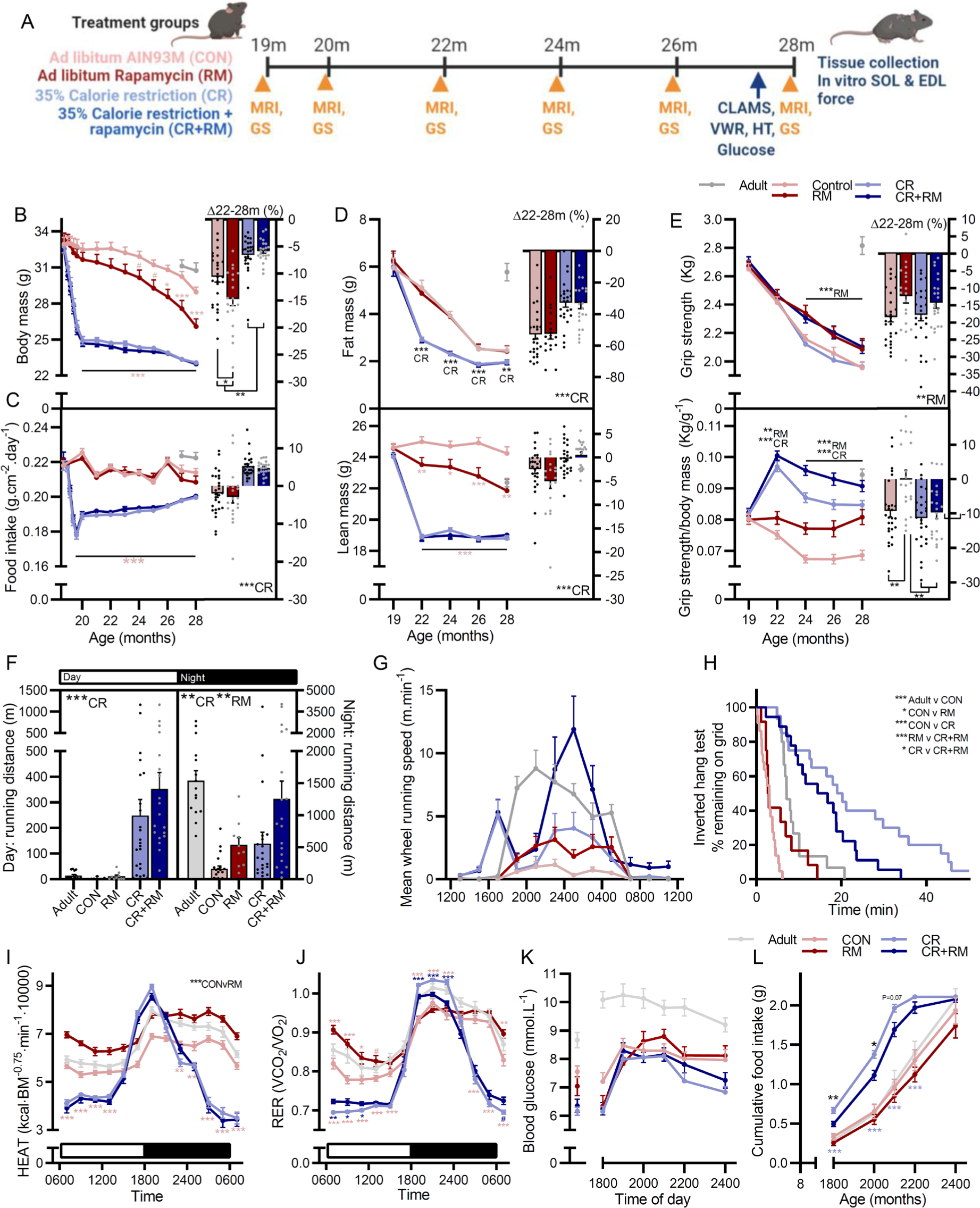
RM and CR exert distinct and additive effects on whole-body muscle function and metabolism. **(A)** Experimental design schematic showing experimental groups as well as time course of physiological measures. Repeated measures (left) of **(B)** body mass, **(C)** food intake normalized to body surface area, **(D)** whole-body fat (upper) and lean (lower) mass as well as **(E)** absolute (upper) and body mass normalized (lower) all-limb grip strength measured across the treatment period from 19 to 28 months as well as the percentage change between 22 months after adaptation to CR and 28 months (right). **(F)** Day and night-time voluntary running distance and **(G)** running speed patterns across a 24 hour period, as well as **(H)** Kaplan–Meier plot for the inverted grip-hang test performed prior to endpoint measures at 28 months. Whole-body metabolic analysis of **(I)** energy expenditure normalized to body surface area and **(J)** respiratory exchange ratio reported every 2 h across one full day (white)/night (black) cycle in the month prior to endpoint measures. **(K)** blood glucose levels and **(L)** voluntary food intake over the night-time feeding period following a day-time fast in 10mCON, 28mCON, 28mRM, 28mCR and 28mCR+RM groups. Data are presented as mean ± SEM. Two-way ANOVAs with Tukey post hoc tests (A–G and I–L) and Mantel–Cox log rank tests (H) were used to compare the data. *, **, and *** denote a significant difference between groups of *P* < 0.05, *P* < 0.01, and *P* < 0.001, respectively. # denotes a trend where 0.05 < *P* < 0.10. Colored asterisks refer to the group of comparison.

Repeated grip strength measurements spanning the treatment period showed a clear RM-induced attenuation in the progressive loss of both absolute and body mass normalized grip strength (Fig. 6E) starting at 24 months of age. CR also induced a strong increase in body mass normalized grip strength between 19 and 22 months, independent of RM treatment. RM and CR also induced distinct changes in voluntary wheel running activity. While CR increased both day and night-time running distance independent of RM treatment, RM increased night-time running distance independent of CR (Fig. 6F-G). CR also potently improved inverted grid-hang time independent of RM treatment (Fig. 6H).

RM and CR individually induced changes in whole-body metabolism consistent with our previous observations (*13*) (Fig. I-J). RM induced a consistent increase in energy expenditure normalized to body surface area in control mice, but not in CR mice (Fig. 6I), consistent with both RM and CR reducing non-functional tissue mass (Fig. S5B-C). In contrast, RM blunted both the CR-induced increase in RER during the first half of the night and the CR-induced decrease in RER during day-time hours (Fig. 6J). Since prolonged RM treatment is known to inhibit mTORC2 and thereby glucose uptake, we wondered whether alterations in blood glucose or feeding behavior may explain the effect of RM on whole-body fuel utilization. Blood glucose was not different between CON and RM or between CR and CR+RM groups at 8 am or at 6 pm after a 10 h fast (Fig. 6K). Likewise, RM did not alter the blood glucose response to the reintroduction of food between 6pm and 12pm (Fig. 6K). CR mice ate their 2.1 g food allocation at a higher rate than *ad libitum*-fed mice (Fig. 6L). Interestingly, CR+RM mice ate their food allocation more slowly than CR mice, which may serve as a strategy to prevent high blood glucose levels in the face of a RM-induced impairment in glucose uptake.

Next we used 2-WAY ANOVAs to determine whether CR and RM also exerted distinct, independent effects on muscle mass. Consistent with improvements in whole-body muscle function, significant CR main effects were observed for relative TA, QUAD, EDL, PLA, GAS, brachioradialis (BR) and biceps brachii (BIC) muscle mass (Fig. 7A). Again, the effects of RM were either distinct or additive to those of CR, with significant RM main effects for TA, QUAD, EDL, PLA, BR, BIC and TRI. As we have previously observed, specific muscles have different responses to specific interventions (*13*). RM increased both relative and absolute muscle mass in the TRI muscle, while CR did not affect relative mass and reduced absolute TRI muscle mass (Fig. 7A and S5A). On the other hand, CR, but not RM improved relative GAS muscle mass, while CR significantly reduced and RM tended to reduce absolute GAS mass. The only interaction effect for RM and CR was observed for the SOL muscle, where RM improved relative muscle mass in CR mice but not *ad libitum* fed mice.

**Figure 7.**
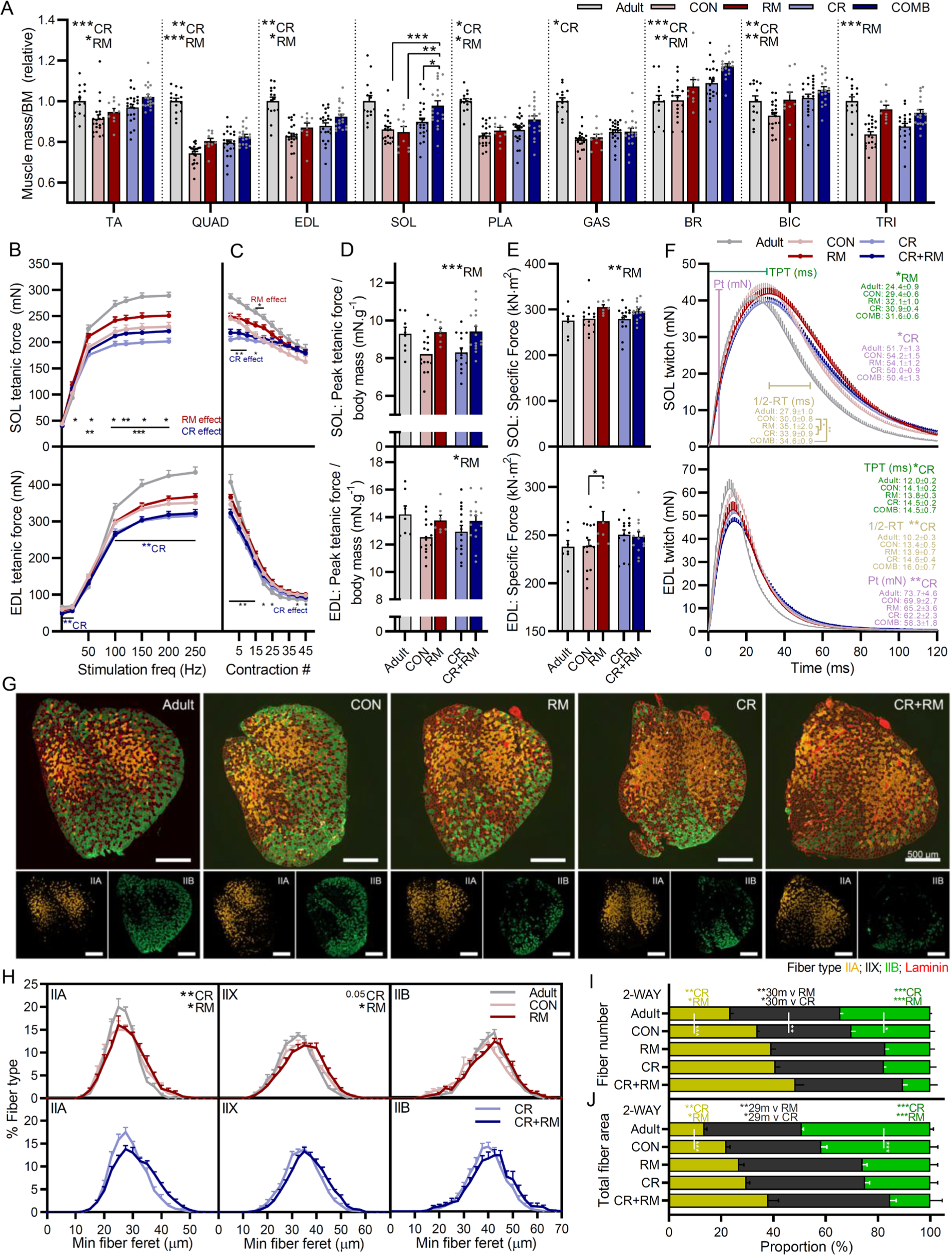
CR and RM have additive effects on muscle mass, function and fiber size and composition. **(A)** Muscle mass for tibialis anterior (TA), *quadriceps* (QUAD), *extensor digitorum longus* (EDL), *soleus* (SOL), *plantaris* (PLA), *gastrocnemius* (GAS), brachioradialis (BR), biceps brachii (BIC) and *triceps brachii* (TRI) were averaged across both limbs, normalized to body mass and then to 10-month-old control mice. Isolated muscle function parameters, including **(B)** force-frequency curve and **(C)** fatigue response to multiple stimulations, **(D)** peak force normalized to body mass, **(E)** peak force normalized to cross sectional area (specific force), and **(F)** mean twitch responses including time-to-peak tension (TPT), half-relaxation time (1/2-RT) and peak twitch (Pt) for SOL (top panel) and EDL muscle (bottom panel). **(G)** Representative cross sectional images along with **(H)** fiber-type specific minimum fiber feret distribution showing *ad libitum*-fed (upper) and calorie restricted (lower) groups, **(I)** fiber type-specific fiber numbers and **(J)** total fiber type-specific cross sectional area of the forelimb muscle brachioradialis (BR) stained with antibodies against type I (blue), type IIA (yellow), and type IIB (green) fibers as well as laminin (red), while fibers without staining were classified as IIX. Data are presented as mean ± SEM. Two-way repeated-measure ANOVAs with Tukey’s post hoc tests were used to compare between data. *, **, and *** denote a significant difference between groups of *P* < 0.05, *P* < 0.01, and *P* < 0.001, respectively. Colored asterisks refer to the group of comparison.

In line with the beneficial effects of RM treatment on whole-body muscle function and muscle size, independent of *ad libitum* or restricted feeding, RM main effects for *in vitro* muscle function properties were observed in the slow-twitch soleus muscle and to a lesser extent the fast-twitch EDL muscle (Fig. 7B-F). CR resulted in significant reductions in absolute tetanic force at multiple stimulation frequencies in the SOL (Fig. 7B, upper) and EDL (Fig. 7B, lower). RM significantly increased absolute tetanic force from a stimulation frequency of 20Hz in SOL, but not EDL muscle. Despite initial contraction forces below that of *ad libitum* fed mice, tetanic force in calorie restricted mice approached that of *ad libitum*-fed mice after 20 contractions in SOL (Fig. 7C, upper) and from 30-35 contractions in EDL (Fig. 7C, lower). RM, but not CR, induced significant increases in peak tetanic force normalized to body mass in both SOL (Fig. 7D, upper) and EDL (Fig. 7D, lower). Rapamycin also induced a significant increase in muscle quality, as evidenced by higher specific forces in SOL muscle (Fig. 7E, upper) independent of feeding status and in EDL muscle (Fig. 7E, lower) from *ad libitum* fed mice. Analysis of muscle twitch properties show that RM slows SOL time-to-peak tension (TPT), while CR slowed EDL TPT (Fig. 7F). RM, CR and CR+RM mice all had longer half-relaxation times in SOL muscle, while CR induced a slowing of half-relaxation time in EDL.

Finally, we have shown that CR promotes a fast-to-slow fiber type transition (Fig. 2G-H) while RM also promotes a fast-to-slow fiber type transition and additionally increases the size of IIA and IIX fibers (*13*). Based on the preferential effect of both CR and RM on slow-twitch fibers and our previous observations that RM exerts negative side effects in some hindlimb muscles that may relate to their susceptibility to denervation, we analyzed the fiber-type composition and fiber cross sectional area in BR muscle, a forelimb muscle involved in grasping and arm flexion that contains a high proportion of IIA and IIX fibers (Fig. 7G-J). In BR muscle, both CR and RM independently increased the size of IIA and IIX fibers, with main effects for both treatments (Fig. 7H). Likewise, both CR and RM showed main effects for higher IIA fiber number and lower IIB fiber number (Fig. 7I), an effect also induced by age alone.

Together, the increase in IIA fiber size and number and the parallel decrease in IIB fiber number observed with age, CR and RM resulted in a change in proportional total fiber area from 13.6, 37.3 and 49.1% for IIA, IIX and IIB fibers in 10-month-old adult mice to 38.1, 46.6 and 15.3% for IIA, IIX and IIB fibers in CR+RM treated mice (Fig. 7J).

Together, these data confirm that both CR and RM treatment exert distinct and often additive effects on whole-body muscle function and metabolism, the size and fiber type composition of specific muscles as well as isolated muscle function properties. Therefore, these findings support the idea that long-term mTORC1 suppression via RM can benefit aging skeletal muscle independent of CR.

## DISCUSSION

The mechanism underlying the potent and robust anti-aging effects of CR have long fascinated the research community. An early theory proposed mTORC1 suppression as a central part of CR-induced longevity (*8*). Research showing long-term RM treatment extended lifespan in mice seemed to confirm this theory (*7*). However, despite both CR and RM blunting mTORC1 activity and promoting autophagy (*18, 19*), their effects on insulin signaling and glucose tolerance diverge (*19, 20*) and molecular profiles following short-term CR and RM in adult mouse liver show predominately distinct signatures (*21, 22*). Our multi-muscle transcriptomic data from 30-month-old mouse muscle support the concept that CR and RM have distinct mechanisms. Importantly, our study compares the effects of CR and RM treatment during the entire period of sarcopenic development (15 to 30 months), therefore representing both signaling responses to CR and RM as well as the cumulative effect of the treatments on age-related signaling. To improve the current understanding of the molecular mechanisms contributing to sarcopenia, we made our data publicly available through the user-friendly web application SarcoAtlas (https://sarcoatlas.scicore.unibas.ch/), building on our previously released sarcopenia data sets (*13, 14*).

The divergent gene expression patterns induced by CR and RM do not necessitate that entirely non-overlapping mechanisms are responsible for the beneficial effects of CR and RM. Rather, while starting from different points, these treatments could converge on the same effector pathways. Here we addressed the question of whether CR-induced mTORC1 suppression is sufficient to make RM redundant and vice versa, and provided definitive evidence that CR and RM exert non-overlapping and often complementary effects in aging mouse skeletal muscle. Specifically, CR improves skeletal muscle function and quality independent of mTORC1 suppression and RM improves skeletal muscle function in CR mice. While mTORC1 inhibition as a strategy to fight aging arose from attempts to pharmacologically reproduce CR, the implications stemming from the apparent invalidation of this assumption are far greater to the aging field. Assuming that a true CR mimetic can eventually be identified, the prospect of multiple, additive interventions to slow aging is an exciting prospect.

While the specific mechanisms responsible for the pro-longevity effects of CR have proved challenging to pin down, many plausible theories have been developed. The evolutionary model of CR proposes that under energy scarcity, an organism actively allocates energy towards somatic maintenance at the expense of reproduction, thereby extending lifespan in the hope that circumstances will sufficiently improve to once again support reproduction (*38, 39*). This theory is supported by observations that CR promotes the expression of maintenance and repair processes (*40, 41*). Signs of a CR-induced promotion of somatic maintenance was also observed in TSCmKO mice in the absence of mTORC1 suppression or reduced ER stress (Fig. 5). A CR-mediated induction of the somatic maintenance genes *Xbp1* and *Keap1* was associated with less P62 accumulation and improved skeletal muscle integrity (Fig. 5), highlighting the fact that skeletal muscle fiber mTORC1 suppression is not solely responsible for the beneficial effects of CR on muscle homeostasis.

The evolutionary model assumes that CR-induced physiological changes are inherently pro-longevity, however, when returned to an energy-rich diet, calorie restricted *Drosophila melanogaster* reproduce less and experience greater mortality than their age-matched, non-restricted counterparts (*2*). Likewise, the beneficial effects of CR are quickly lost upon refeeding in rodents (*42*) and in humans, metabolic adaptions to CR promote strong weight gain upon refeeding (*43*). Therefore, rather than actively promoting longevity, CR may provide an escape from the cost of an energy-rich diet (*2*). The fact that sedentary, *ad libitum*-fed mice often become obese in aging studies ^12^ and that the propensity of mouse strains to become obese strongly correlates with the longevity effects of CR (*44*) support this theory. Importantly, in our study we used a combination of an AIN-93 maintenance diet and a controlled feeding regime to limit the food intake of *ad libitum*-fed mice with a penchant for overeating (Fig. 1), thereby limiting the potential negative effects of an obese and unhealthy control group, a common criticism of CR experiments (*44*).

An alternate theory proposes that rather than an evolved, programmed tradeoff between somatic maintenance and reproduction, the pro-longevity effects of CR may simply result from a passive, serendipitous response to metabolic stress (*2*) such as a lower metabolic rate (*45*), reduced mTORC1 activity (*8*) or hormetic response (*46*). In line with such an interpretation, long-term CR did not slow the absolute loss of all-limb grip strength nor did it reverse age-related gene expression changes. However, the CR-induced alterations in body composition led to marked improvements in body mass-normalized grip strength, which proceeded to decline with age in line with *ad libitum*-fed mice, but from a new higher starting point (Fig. 1 and 6). Irrespective of the permanence of CR-induced longevity and whether the effects are actively or passively conferred, the strong link between CR and beneficial health and lifespan outcomes means that closely examining the molecular and physiological responses to CR is a promising avenue to identify pathways and phenotypes that can be exploited to improve muscle function and consequently, the quality of life of aging individuals. The same is true for the pro-longevity and anti-sarcopenic effects of rapamycin. While our rationale for comparing these two interventions originally centered on identifying a core set of pathways regulated by aging and commonly counter-regulated by CR and RM, our data show that CR and RM induce strikingly distinct gene expression profiles, with those genes and pathways that do overlap being regulated in predominately opposing directions (Fig. 3).

For example, CR specifically increases genes involved in lipid metabolism and RM specifically suppresses genes involved in insulin signaling, while the strongest counter-regulated gene expression cluster maps to ECM remodeling, including many collagen genes, which are reduced in aging muscle, restored by rapamycin but further decreased by CR. ECM gene expression coincides with changes in muscle size, increases during muscle hypertrophy (*47*), and decreases during experimental muscle atrophy (*48*) as well as sarcopenia in rodents (*13, 14, 49, 50*). Declining collagen expression has also been linked to organismal aging, while autophagy-inducing interventions restore collagen expression and promote longevity (*51, 52*). While this opposing action of CR and RM on ECM gene expression could be seen as counteractive, it may also represent different means of addressing the same perturbation. That is, age-related stressors can be counteracted by either blunting the perturbation or enhancing the inherent coping mechanisms. Indeed, gene expression profiling studies show that changes in gene expression resulting from mutations in mice that shorten lifespan positively correlate with mutations and interventions that extend lifespan (*53*). In other words, organisms naturally boost stress responses to cope with life-shortening perturbations, while interventions that moderately stimulate these stress responses, such as CR, extend lifespan. Alternatively, changes in ECM gene expression may simply reflect changes in protein turnover or absolute muscle size induced by the treatment. In line with this hypothesis, CR blunts and RM boosts polysome loading, an indicator of protein turnover, in skeletal muscle (*22*). The complexity of biological responses to aging and treatments such as CR and RM make interpreting directionality of gene expression changes inherently challenging. It is therefore imperative that molecular profiles be accompanied by thorough phenotypic characterizations.

Perhaps the most strikingly consistent muscle phenotype displayed by CR mice was a fatigue resistant, fast-to-slow muscle fiber property switch, including an increase in number and proportional cross sectional area covered by slower-type fibers across five different muscles as well as slower twitch properties (Fig. 2 and 7) and higher whole-body relative muscle endurance (Fig. 1 & 6). A similar, although less pronounced phenotype was also observed in response to long-term RM treatment (*13*) (Fig. 7). Despite this seemingly overlapping effect of the two treatments, the CR- and RM-induced fast-to-slow fiber type conversion was additive in the forelimb *brachioradialis* muscle of CR+RM treated mice (Fig. 7). Since slower-type fibers are more resistant to age-related atrophy, the CR-induced fast-to-slow fiber-type transition, despite the absolute decrease in muscle mass, may ultimately preserve muscle fiber number and function in aging mice, as previously observed in muscle-specific BDNF knockout mice, which display a fast-to-slow fiber type transition and increased resistance to age-related muscle loss(*31*).

A specific feature of CR believed to contribute to its beneficial effects in rodent skeletal muscle (*17*) is an increase in physical activity prior to feeding, and may at least partially explain the fast-to-slow fiber transition. This anticipatory behavior, reminiscent of exercise training, is not seen in *ad libitum*-fed mice and was not altered by RM in the current study (Fig. 6). Although the CR-induced quasi exercise training is an artefact of experimental circumstance not applicable to humans, the combined impact of CR, including any benefits afforded through behavioral changes were on top of those induced by RM, particularly with regard to slowing of muscle fiber properties, relative grip strength and voluntary running distance (Fig. 6 & 7). Since muscle adaptations to exercise are accentuated by protein intake immediately before, during or after exercise, the close temporal restriction of activity and food intake in CR mice may be advantageous for maintaining muscle function with age.

While together, our experiments clearly demonstrate that the beneficial effects of CR on muscle do not require mTORC1 suppression and likewise RM treatment is not redundant in restricted mice, the complementary mechanisms allowing the effects of these interventions to compound are less clear. Both CR and RM are well-known to reduce mTORC1 activity, but unlike in RM-treated mice, mTORC1 activity still responds to food intake in CR mice, although its activity is more temporally restricted, and decreases more during fasting periods than in *ad libitum*-fed mice (*54*). Since transient mTORC1 activity remains important in specific tissues, most blatantly highlighted by the severe testicular degeneration experienced by RM-treated mice, it might be expected that CR would represent a more favorable intervention to counter the detrimentally hyperactive mTORC1 frequently observed in aging tissue. Strikingly, our data point in the opposite direction, suggesting that CR-induced mTORC1 suppression is not sufficient to confer all the beneficial effects of RM treatment. Of course, whether this effect relates directly to muscle tissue mTORC1 suppression or via secondary effects of non-muscle tissue mTORC1 suppression is unclear.

Another explanation for the additive effects of CR and RM, could be co-compensation of treatment-specific side effects. A prime example relates to glucose tolerance, which is markedly improved in CR mice (Fig. 1), and impaired by RM (*55*). However, blood glucose levels were not habitually high in RM or CR+RM groups, instead, CR+RM-treated mice appeared to curb the ravenous consumption of daily food allocation seen in CR mice, thereby distributing energy availability over a prolonged period (Fig. 6K-L). Metabolic analyses showed that this subdued rate of food intake coincided with a lower RER during the early night-time eating phase but also a higher RER during the early day-time inactive period in CR+RM compared to CR mice (Fig. 6J). Whether this smoothing of diurnal fluctuations in energy utilization affords benefits for CR+RM mice remains to be tested. Along a similar line, we previously reported that specific muscles respond differently to rapamycin, an effect correlated with the presence (muscle not protected by RM, i.e. GAS) or absence (muscle protected by RM, i.e. TRI and TA) of pro-sarcopenic side effects (*13*). Although a CR-induced amelioration of these negative effects of RM could explain the combined effects, the same muscle-specific effects of RM were observed in CR+RM mice, with less age-related muscle loss in the TRI and TA, but not in the GAS (Fig. 7A), hinting that further attenuation in muscle aging may be possible beyond combined CR and RM.

Together, our results conclusively demonstrate that CR and RM exert distinct, non-overlapping and frequently additive effects in aging skeletal muscle. The striking failure of RM to recapitulate the effects of CR and more surprisingly, the failure of CR to recapitulate the effects of RM raises the exciting prospect of multiple, additive interventions to counteract sarcopenia. Further work is needed to systematically isolate targetable processes and examine the resulting interactions in order to develop optimal strategies to counteract sarcopenia and promote healthy aging.

## METHODS

### Animal care

All procedures were performed in accordance with Swiss regulations for animal experimentation and approved by the veterinary commission of the Canton Basel-Stadt. Male, C57BL/6JRj mice for aging studies were purchased from the aging colony at Janvier Labs (Le Genest-Saint-Isle, France). Transgenic TSCmKO mice and their genotyping were previously described (*33, 56*). Littermates floxed for *Tsc1* but not expressing HSA-Cre-recombinase were used as controls. For all studies, mice were kept in single cages under a fixed 12h light-dark cycle (6 am to 6 pm) at 22°C (range 20-24°C) and 55% (range 45-65%) relative humidity and were acclimatized to individual housing and the control diet for 3-4 weeks before the start of the experiment.

### Aging studies

In the first aging study (results presented in Figures 1-3), two independent groups of mice were calorie restricted starting at 20 (n=11) or 15 (n=21) months of age. 10mCON, 30mCON and 30mRM data were generated alongside 30mCR, but data from these groups have largely been previously reported (*13*). For appropriate interpretation of the previously unreported CR data, 10mCON and 30mCON data have been included. For combined CR+RM experiments (Figures 6 and 7), all groups and data were newly generated. After 1-month acclimatization to individual housing and the standardized AIN-93M diet (TestDiet, 58M1-9GH6) containing 488 ppm Eudragit (Emtora), food intake of CR mice was incrementally reduced to 90%, 80% and 70% for 1 week each before being maintained at 65% of mean baseline food intake. To avoid malnutrition, the concentration of vitamins and minerals were increased 1.43 × in CR mice (58M1-9GH8). Rapamycin treated *ad libitum*-fed and CR mice received an AIN-93M diet containing 42 or 48 mg·kg^-1^ active encapsulated (Eudragit) rapamycin (TestDiet, 58M1-9GH7), respectively. The concentration of RM was increased in CR+RM mice in an attempt to counter the discrepancy between body weight (∼-22%) and food intake reductions (-35%) and deliver a similar dose of rapamycin per kg body weight. In the week before starting the experiment, food intake, body mass and composition (via EchoMRI) and grip strength were measured and used to balance group selection. Each group (CON, CR, RM and CR+RM) contained mice with an almost identical mean and standard deviation for each measurement within experimental groups. Food intake and body mass were measured weekly. As previously described (*13*), weight gain and obesity was limited in *ad libitum*-fed mice via daily food restriction to that of the control group mean (3.1 g) in mice with a propensity for overeating and weight gain.

### TSCmKO studies

Mice were progressively adapted to calorie restriction by incremental reductions in food intake of 10% per week starting at 12 weeks of age, with mice receiving 70% of the genotype mean at the start of the 3^rd^ week. Baseline food intake was determined over a 3-week period prior to CR. *Ad libitum*-fed WT and TSCmKO mice ate 3.05 and 3.45 g·day^-1^ food, respectively, under baseline conditions. After CR habitualisation, WT-CR and TSC-CR groups were therefore given 2.1 and 2.4 g·day^-1^ food for 6 months. These experiments were performed over three separate trials using both female and male mice and either an AIN-93M standardized diet (D10012M; KLIBA NAFAG) or a standard chow diet (KLIBA NAFAG-3432). Since results were comparable across all trials, food and sexes, data were pooled for analysis.

### Body composition analysis

Fat mass and lean mass were measured on restrained, conscious mice using an EchoMRI-100 (EchoMRI Medical Systems).

### Whole-body muscle function

Mice were initially familiarized to voluntary running wheels over a 48-72 h period. Mice were then given free access to voluntary running wheels for a 24 h period every two months in the first study and once for a 48 h period in the subsequent CR+RM study, with data for the final 24 h used for analysis. Inverted grid hang time was measured by placing mice on a wire grid, which was slowly turned upside down and positioned over a ∼40 cm high box containing a foam pad at the bottom. Performance was taken as the longest time a mouse could hold onto the grip across three trials separated by at least 30 min. All-limb grip strength was measured using a small grid attached to a force meter (Columbus Instruments). Mice firmly holding the grid with all four paws were gently pulled horizontally at a consistent speed until the grasp was broken. Performance was taken as the median of 3-5 trials with at least 10 min rest between tests. Trials in which the mouse actively pulled on the grid while the test was underway where discarded. The same researcher performed all grip strength measurements at a similar time of day.

### Comprehensive laboratory animal monitoring system (CLAMS)

CLAMS (Columbus Instruments, Columbus, OH) was used to measure whole-body metabolic parameters including energy expenditure, oxygen consumption (VO_2_), CO_2_ production (VCO_2_) and the ratio of VCO_2_ to VO_2_ or respiratory exchange ratio (RER), as well as locomotory activity. RER is dependent on energy substrate utilization, ranging from above 1 to 0.7, which indicates preferential use of carbohydrates and lipids, respectively. Locomotor activity was measured on X, Y and Z axes using infrared beams. Energy expenditure was calculated using VO_2_ and RER values and subsequently normalized to body surface area (body mass^-0.75^) to account for pronounced changes in body size associated with CR. Data were collected for three consecutive days, with the final 24 h period (6 am to pm) used for analysis.

### In vitro muscle force

*In vitro* muscle force was measured in the fast-twitch *extensor digitorum longus* (EDL) and slow-twitch *soleus* muscles. After careful isolation, muscle tendons were tied with surgical suture at each end and mounted on the 1200A Isolated Muscle System (Aurora Scientific, Aurora, ON, Canada) in an organ bath containing 60 mL of Ringer solution (137 mM NaCl, 24 mM NaHCO_3_, 11 mM Glucose, 5 mM KCl, 2 mM CaCl_2_, 1 mM MgSO_4_, 1 mM NaH_2_PO_4_) gassed with 95% O_2_; 5% CO_2_ at 30 °C. After defining optimal length, muscles were stimulated with 15 V pulses. Muscle force was recorded in response to 500 ms pulses of 10-250 Hz. Muscle fatigue was assessed by 6 min of repeated tetanic stimulations at 200 Hz for EDL and 120 Hz for SOL, respectively, separated by 8 sec.

### Immunostaining of muscle cross sections

Muscles were mounted at resting length in optimal cutting temperature medium (O.C.T, Tissue-Tek) and snap-frozen in thawing isopentane for ∼1 min before transfer to liquid nitrogen and storage at -80°C. Muscle sections (10 µm) were cut from the mid belly at -20° C on a cryostat (Leica, CM1950), collected on SuperFrost Plus (VWR) adhesion slides and stored at -80° C. Sections from each experimental condition were always mounted on the same slide to ensure accurate comparisons. For fiber typing, sections were blocked and permeabilized in PBS containing 10% goat serum and 0.4% triton X-100 for 30 min before being incubated for 2 h at RT in a primary antibody solution containing BA-D5, SC-71, BF-F3 which were developed by Prof. Stefano. Schiaffino and obtained from the Developmental Studies Hybridoma Bank developed under the auspices of the National Institute of Child Health and Human Development and maintained by the University of Iowa Department of Biology as well as laminin (#11575; Abcam) and 10% goat serum. After incubation in primary antibodies, sections were washed 4 × 10 min in PBS and then incubated in a secondary antibody solution containing DyLight 405 (#115-475-207, Jackson), Alexa568 (#A-21124, Invitrogen), Alexa488 (#A-21042, Invitrogen), Alexa647 (#711-605-152, Jackson) and 10% goat serum. Sections were then washed 4 × 10 min in PBS and mounted with ProLong™ Gold antifade (Invitrogen). Muscle sections were imaged at the Biozentrum Imaging Core Facility with an Axio Scan.Z1 Slide Scanner (Zeiss) equipped with appropriate band-pass filters. Fiji macros were developed inhouse to allow an automated analysis of muscle fiber types (based on intensity thresholds) and muscle cross-sectional area (i.e., minimal Feret’s diameter; based on cell segmentation) (*31*). All macros and scripts used in this study are available upon request. For P62 staining, TA sections were thawed for 10 min at RT, fixed in 4% PFA for 6min, neutralized in 0.1M Glycine (pH7.4) at RT for 2 × 15 min and then blocked at RT for 1.5 h in blocking solution containing 3% IgG free Bovine Serum Albumin (BSA), 1% Fab anti-mouse IgG (Jackson) and 0.25% Triton-X. Slides were then incubated in primary antibody solution containing P62 (GP62-C, 1:300), laminin (ab11576, Jackson 1:300) and 3% BSA overnight at 4°C. The next day, slides were washed 3 × 10 min in PBS and incubated in secondary antibody solution containing DaGP Cy3 (706-165-148; Jackson; 1:500) and GaRt 488 (112-545-003, Jackson; 1:500) for 1.5h at RT. Afterwards, sections were washed 2 × 10 min in PBS and mounted with Vectashield DAPI (Vector Laboratories).

### Protein extraction and Western blot analysis

For TSC-CR studies, TA and GAS muscles were snap-frozen, pulverized in liquid nitrogen and lysed in RIPA buffer before sonication and 2 h incubation at 4° C. Lysates were then centrifuged at 16,000 *g* for 30 min at 4° C to remove insoluble material. Protein concentration was measured (BCA assay) and normalized with RIPA buffer before being heated for 5 min at 95° C in Laemmli buffer (0.1 M Tris-HCl pH 6.8, 10% Glycerol, 2% SDS, 0.04% Bromphenolblue, 1% β-mercaptoethanol). Proteins were separated on 4-12% Bis-Tris protein gels (NuPAGE, Life Technologies) and transferred onto nitrocellulose membranes. Membranes were blocked for 1 h at RT in 3% BSA in TBS containing 0.1% Tween20 and incubated overnight with primary antibodies in blocking solution at 4° C. The next day, membranes were washed 3 × 10 min in TBS before incubation in secondary horseradish peroxidase-conjugated antibodies, diluted in blocking solution. Membranes were then washed 3 × 10 min in TBS before immunoreactivity was visualized using the KLP LumiGlo Chemiluminescence Substrate Kit (Seracare) with a Fusion Fx machine (Vilber). Protein abundance was quantified using FusionCapt Advance (Vilber) as mean grey value minus background and then normalized to a housekeeping protein. Western blots for all proteins were performed on TA muscle. For key proteins (phospho and total mTOR, S6, 4EBP1 and AKT, as well as P62 and LC3B), western blots were also performed on GAS muscle. Results in both muscles were comparable and were subsequently averaged for each mouse for further analysis. All primary antibodies were from Cell Signaling Technology: p^s2448^mTOR (#2971), mTOR (#2972), p^S240/244^S6 (#5364), p^S235/236^S6 (#2211), S6 (#2217), p^S65^4EBP1 (#9451), 4EBP1 (#9452), LC3B (#2775), p^S757^ULK (#6888), ULK (#8054), Beclin1 (#3495), Bnip3 (#3769), p^T246^PRAS40 (#2997), PRAS40 (#2610), p^S473^AKT (#4058), AKT (#9272) and BiP (#3177) at a dilution of 1:1000, except ATF4 (sc-200, Santa Cruz, 1:1000), FGF21 (AF3057, 1:500), α-actinin (A7732, Sigma, 1:5000) and P62 (GP62-C, Pro-Gene, 1:1000).

### RT-qPCR

Snap frozen gastrocnemius muscles were pulverized and lysed in RLT buffer (Qiagen). RNA was extracted using the RNeasy® Mini Kit (Qiagen), with Proteinase K and DNase treatment, according to the supplier’s instructions. RNA purity was determined using a Nanodrop ONEC (Thermo Scientific). cDNA was generated with the iScript^TM^ cDNA Synthesis Kit (Bio-Rad) using 500 ng of extracted RNA according to supplier’s manual. cDNA samples were stored at -20°C. RT-qPCR was performed in duplicate with the LightCycler 480 (Roche Diagnostics) instrument using LightCycler 384-well plates with sealing foil (Roche). The reaction volume of 10 μl contained, FastStart Essential DNA Green Master Mix (2X, Roche), forward and reverse primers and cDNA template (1:5 diluted). Primers were designed using Genious®10 software (*57*) and specificity confirmed by the Basic Local Alignment Search Tool (BLAST) (*58*). Potential hairpin formation, complementarity and self-annealing sites were verified to be negative by OligoCalc (*59*). The amplification of a single PCR product was confirmed with a melting-point dissociation curve and raw quantification cycle (Cq) values were calculated by a LightCycler 480. Data were analyzed using the comparative Cq method (2^−ΔΔCq^). Raw Cq values of target genes were normalized to Cq values of a housekeeping gene (β-actin), which was stable between conditions, and then further normalized to the control group for ease of visualization. Primers used are as follows:

*Map1lc3a: Fwd-*GTTGGATGTGTTCTGTCTCGTCAC; *Rev*-CTACGTGATTATTTCCGTGTTGCT

*Atg7: Fwd*-TGCAGTTCGCCCCCTTTAAT; *Rev*-CAGGCGGTACTCGTTCAACT

*Bnip3: Fwd-*TTCCACTAGCACCTTCTGATGA; *Rev*-GAACACCGCATTTACAGAACAA *Gabarapl1: Fwd-*CATCGTGGAGAAGGCTCCTA; *Rev*-ATACAGCTGGCCCATGGTAG

*Sqstm1: Fwd-*TACTCGAACGACACAAGGGA; *Rev*-GACTCAGCTGTAGGGCAAGG

*Ctsl: Fwd-*GTGGACTGTTCTCACGCTCA; *Rev*-TCCGTCCTTCGCTTCATAGG

*Trib3: Fwd-*GGACAAGATGCGAGCCACAT; *Rev*-CCACAGCAGGTGACAAGTCT

*Xbp1: Fwd-*TGGCCGGGTCTGCTGAGTCCG; *Rev*-GTCCATGGGAAGATGTTCTGG

*Keap1: Fwd-*GGCAGGACCAGTTGAACAGT; *Rev*-GGGTCACCTCACTCCAGGTA

*Gsta1: Fwd-*CCAGAGCCATTCTCAACTA; *Rev*-TGCCCAATCATTTCAGTCAG

### RNA extraction for aging-calorie restriction data sets

For the aging-CR-RM data set, RNA extraction was performed as previously described in detail (*13*). Briefly, TA, TRI, GAS and SOL muscles from 6 mice per group were pulverized and lysed in RLT buffer (Qiagen) and treated with proteinase K (Qiagen) and DNAse. RNA was extracted with an iColumn 24 (AccuBioMed) with an AccuPure Tissue RNA Mini Kit (AccuBioMed). RNA purity and integrity was examined with a Bioanalyser (Agilent). RNA concentration was determined with a Quant-iT™ RiboGreen™ RNA assay kit and Qubit flurometer (Invitrogen). Libraries were prepared with TruSeq Stranded mRNA HT Sample Prep Kit. Stranded, paired-end sequencing with 101 base pair read length was performed on an Illumina HiSeq2500 platform. A single outlier in the 30mCON SOL group was identified and removed from further analysis based on a clear technical error.

### Statistical analysis

All values are expressed as mean ± SEM unless stated otherwise. Data were tested for normality and homogeneity of variance using a Shapiro-Wilk and Levene’s test, respectively. Data were analysed in GraphPad Prism 8. Student t-tests were used for pairwise comparisons, while one-way ANOVAs with Fisher’s LSD post hoc tests were used to compare between three groups, so long as the ANOVA reached statistical significance. Two-way ANOVAs or two-way repeated measures ANOVAs for multiple recordings over time, with Sidak or Tukeys post hoc tests were used to compare between groups with two independent variables. Both significant differences (P < 0.05) and trends (P < 0.1) are reported where appropriate.

### RNA-Seq data processing

Paired-end RNA-Seq reads were subjected to 3’ adapter (mate 1 5’-AGATCGGAAGAGCACACGTC-3’, mate 2 5’-AGATCGGAAGAGCGTCGTGT-3’) and poly(A)/poly(T) trimming using Cutadapt v1.9.1 (*60*). Reads shorter than 30 nucleotides were discarded. As the reference transcriptome, we considered sequences of protein-coding transcripts with support level 1-3 based on the genome assembly GRCm38 (release 92) for mouse and corresponding transcript annotations from Ensembl database (*61*). The kallisto v0.43.1 software was used to assign filtered reads to mouse transcriptome (*62*). The default options of kallisto were utilized for building the transcriptome index. For aligning stranded RNA-Seq reads, where mates 1 and 2 originated from antisense and sense strands, respectively, the option ‘--rf-stranded’ was used. The option ‘--pseudobam’ was used to save kallisto pseudoalignments to a BAM file.

Mapped reads were then assigned to transcripts in a weighted manner: if a read was uniquely mapped to a transcript, then the transcript’s read count was incremented by 1; if a read was mapped to *n* different transcripts, each transcript’s read count was incremented by 1/*n*. Trimming 3’ adapters and poly(A)/poly(T) stretches, indexing reference transcriptomes, mapping the RNA-Seq reads to transcripts and counting reads assigned to individual transcripts were performed with a Snakemake framework (*63*).

The expression of each transcript *t_i_* was then estimated in units of transcripts per million (TPM) by dividing the read count *c_i_* corresponding to the transcript by the transcript length *l_i_* and normalizing to the library size:

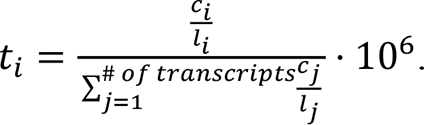

The expression level of a gene was calculated as the sum of normalized expression levels of transcripts associated with the gene. For every gene, read counts of transcripts associated with this gene were also summed up.

### Calculating log-fold changes in the gene expression across conditions

Calculating log-fold changes in the gene expression across conditions was performed with EdgeR available through the R/Bioconductor package (*64*). A gene was included in the analysis only if it had at least 1 count per million (CPM) in the number of samples corresponding to the minimum number of replicates of the same condition across conditions. Obtained log-fold changes were subjected to the hierarchical clustering based on Euclidean distance.

### Aligning gene expression with principal components

The gene expression matrix with samples as columns and log2-transformed gene expression in TPM units as rows was mean centered to make the data comparable both across samples and genes. The centered gene expression matrix was further subjected to the principal component analysis (PCA). First four principal components, PC1, PC2, PC3 and PC4, were defined for each data subset, respectively. Then for each data subset we quantified how much individual genes contributed to the corresponding principal component (*13*). Shortly, we represented genes in the multi-dimensional sample space, localized principal components and calculated projections of vectors associated with genes on principal components and correlations between gene vectors and principal components. We considered a projection of the gene vector on PC significant if its absolute *z*-score value was ≥1.96. The correlation between a gene vector and PC was considered as significant if the absolute value was ≥0.4.

### Gene Set Enrichment Analysis

The distribution of gene sets in ranked gene lists was examined using GSEA(*65*). Ranking was based on log-fold changes in the gene expression between two conditions of interest. Enrichment was considered significant if FDR was less than 0.01.

### Hierarchical Clustering

Hierarchical clustering of genes was based on Euclidean distance between changes in the gene expression in notified conditions (see Fig. 3E).

### Gene ontology analysis

To annotate genes aligned with PCs, we performed the gene ontology (GO) analysis using Database for Annotation, Visualization and Integrated Discovery (DAVID) (*66*) through the R/Bioconductor package called ‘RDAVIDWebService’ (*67*). ‘GOTERM_BP_DIRECT’, ‘GOTERM_MF_DIRECT’ and ‘GOTERM_CC_DIRECT’ categories were used for gene annotation. Background genes for calculating enrichment statistics consisted of all genes expressed in muscle samples (see the section ‘Aligning gene expression with principal components’). GO terms with a p-value less than 0.01 were considered significantly enriched.

### Shiny application

To make CR high-throughput data set and data analysis tools available for the research community, we included them in the previously developed interactive web application ‘SarcoAtlas’ based on the R package Shiny (version 0.14.2, https://cran.r-project.org/web/packages/shiny/index.html). The application supports gene expression plotting, differential expression analysis, principal component analysis and aligning gene expression with principal components. Moreover, the application can submit genes resulting from the analysis to STRING (*68*) to further investigate protein-protein interactions and perform GO analysis. The application can be accessed through the following link: https://sarcoatlas.scicore.unibas.ch/ [10].

## Acknowledgements

We gratefully acknowledge Dr. Mikhail Pachkov for help with data processing, Dr. Alexander Kanitz for testing and Pablo Escobar López for publishing SarcoAtlas and Dr. Erik van Nimwegen for fruitful discussions about data analysis. We also acknowledge the support of the University of Basel’s Quantitative Genomics Facility, in particular Phillippe Demougin for assistance with mRNA-seq sample preparation; the Image Core Facility, in particular Kai Schleicher and; the scientific computing center, sciCORE (http://scicore.unibas.ch/), where calculations were performed.

## Author contributions

Conceptualization: DJH, NM, MAR, MZ, AB, LAT Methodology: DJH, NM, KC, SL, LAT, AB

Investigation: DJH, NM, KC, SL, ABL, ASH, RF, MT, JD, LAT, DB

Data curation and formal analysis: AB, DJH Statistical analysis: DJH, AB

Software: AB Visualization: DJH, AB Supervision: MAR, MZ

Writing-original draft: DJH, AB, MAR

Writing-review and editing. DJH, MAR, NM, AB, MZ, LT Funding Acquisition, MAR, MZ, NM, CH, MS

## Data and materials availability

RNA-Seq data set on four muscles (GAS, TA, TRI and SOL) from six 30-month-old calorie restricted mice was deposited to Gene Expression Omnibus (GEO, https://www.ncbi.nlm.nih.gov/geo/) (*69*) under the accession number GSE171322. RNA-Seq data describing profiles of four muscles (GAS, TA, TRI and SOL) of 10-month-old, 30-month-old and 30-month-old mice treated with rapamycin were previously published (*13*) and available at GEO under the accession number GSE139204. These data are also accessible using the web-based application, SarcoAtlas (https://sarcoatlas.scicore.unibas.ch/). Code is available upon request.

