## Supplementary material for "Distinct and additive effects of calorie restriction and rapamycin in aging skeletal muscle"

1 **Figure S1**

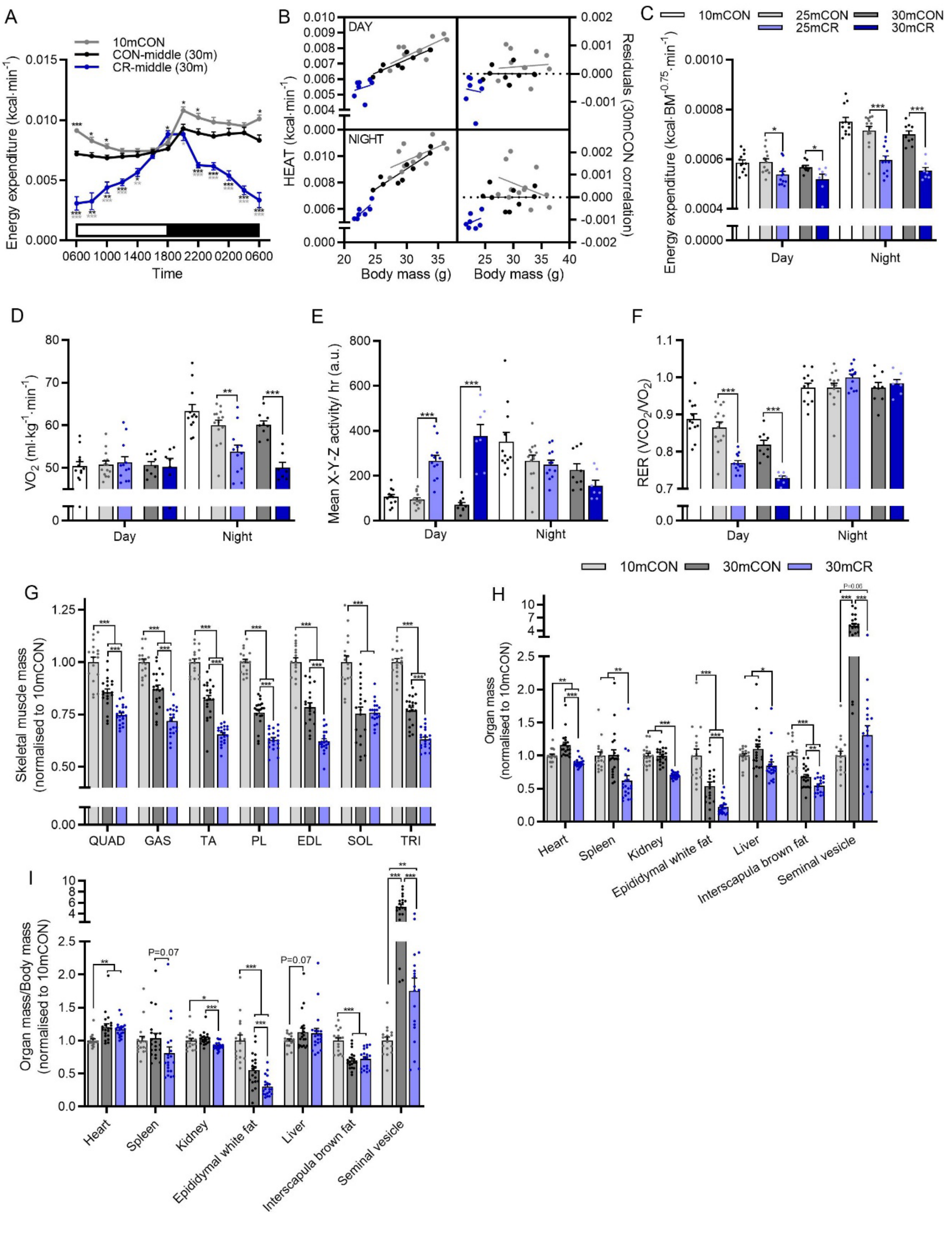

A
10mCON
CON-middle (30m)
CR-middle (30m)
Energy expenditure (kcal·min-1)
Time
B
DAY
NIGHT
HEAT (kcal·min-1)
Residuals (30mCON correlation)
Body mass (g)
C
10mCON
25mCON
25mCR
30mCON
30mCR
Energy expenditure (kcal·BM-0.75·min-1)
Day
Night
D
VO2 (ml·kg-1·min-1)
E
Mean X-Y-Z activity/ hr (a.u.)
F
RER (VCO2/VO2)
10mCON
30mCON
30mCR
G
Skeletal muscle mass (normalised to 10mCON)
QUAD
GAS
TA
PL
EDL
SOL
TRI
H
Organ mass (normalised to 10mCON)
Heart
Spleen
Kidney
Epididymal white fat
Liver
Interscapula brown fat
Seminal vesicle
I
Organ mass/Body mass (normalised to 10mCON)
P=0.07


2

**Figure S1: Metabolic analyses and tissue mass.** (A) Whole-body metabolic analysis of energy expenditure in kcal·min$^{-1}$ reported every 2 h across one full day (white)/night (black) cycle and (B) the relationship between body mass and day or nighttime energy expenditure for 10mCON, 30mCON and 30mCR. Mean day and night whole-body metabolic analysis of (C) energy expenditure normalized to body surface area, (D) VO2 ml·kg$^{-1}$·min$^{-1}$, (E) X-Y-Z activity per hour and (F) Respiratory exchange ratio at 25 and 30-months of age for 30mCON (n=14 at 25m and 9 at 30m) and 30mCR (n= 12 at 25m and 7 at 30m) groups as well as for 10mCON (n=12) mice. (G) Skeletal muscle mass normalized to 10mCON and (H) organ mass normalized to 10m control alone or (I) to body mass and then 10mCON. Group numbers for (G-I) are 17 (10m), 20 (30mCON) and 20 (30mCR) except for brown fat and seminal vesicles (n=19). Data are presented as mean ± SEM. Two-way repeated-measure ANOVA with Sidak or Tukey post hoc tests (A–F), and one-way ANOVA with Fisher's LSD post hoc tests (G-I) were used to compare the data. *, **, and *** denote a significant difference between groups of $P < 0.05$, $P < 0.01$, and $P < 0.001$, respectively. # denotes a trend where $0.05 < P < 0.10$. Colored asterisks refer to the group of comparison.

**Figure S2**

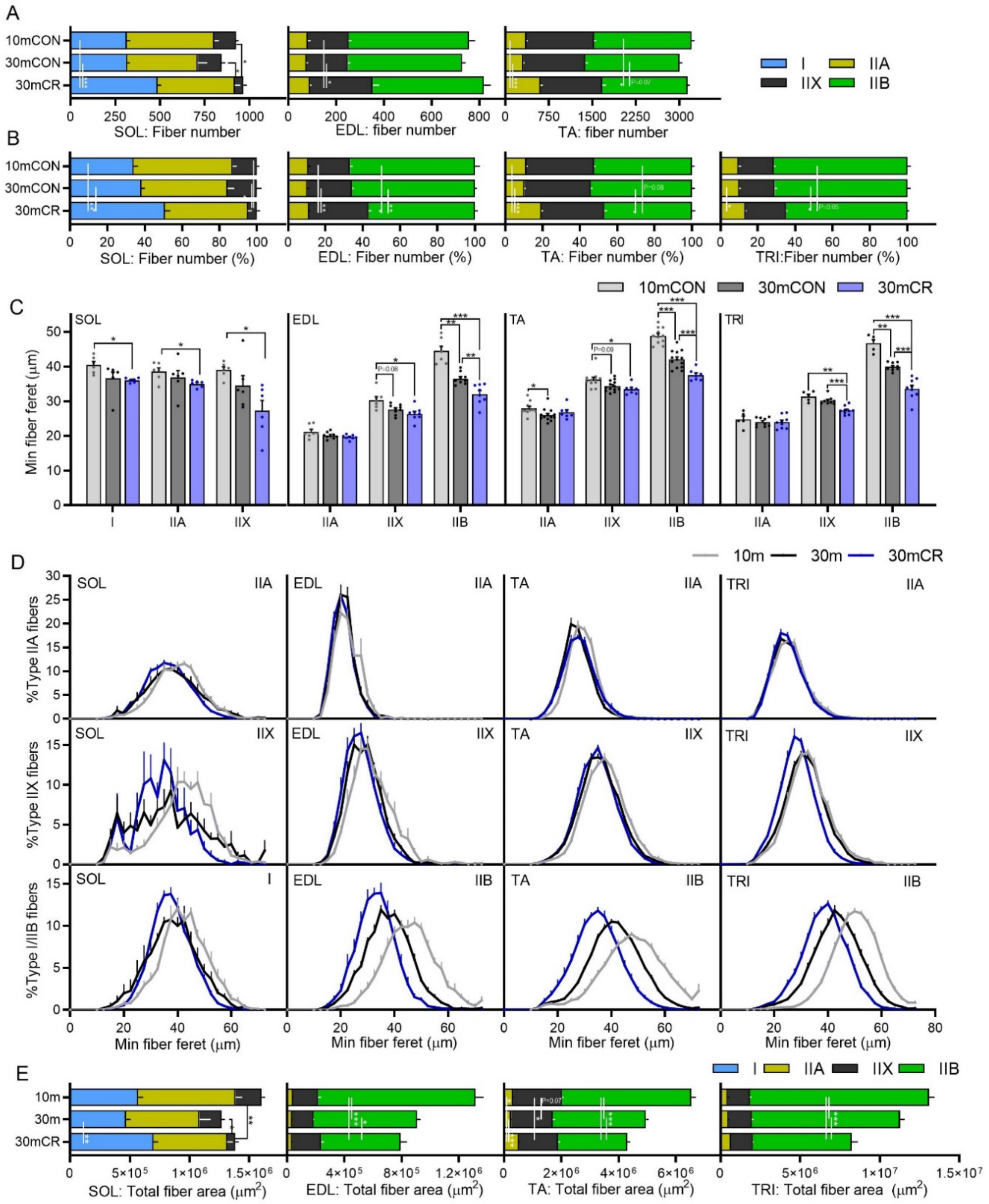


**Figure S2: CR promotes a fast-to-slow muscle fiber phenotype shift.** Fiber-type-specific (A) absolute fiber counts, (B) fiber type proportions, (C) mean minimum fiber feret, fiber size distribution and (E) total fiber cross sectional area on whole cross sections from soleus (SOL; n=6), extensor digitorum longus (EDL; n=7, 9 and 8), tibialis anterior (TA; n=11, 13 and 7) and triceps brachii (TRI; n=5, 9 and 9) for 10mCON, 30mCON and 30mCR muscles stained with antibodies against type I, type IIA and type IIB fibers as well as laminin, while fibers without staining were classified as IIX. Data are presented as mean ± SEM. One-way (D) or two-way repeated-measure (A-C, E) ANOVAs with Fisher's LSD or Tukey's post hoc tests, respectively, were used to compare between data. *, **, and *** denote a significant difference between groups of $P < 0.05$, $P < 0.01$, and $P < 0.001$, respectively. Colored asterisks refer to the group of comparison.

### Figure S3

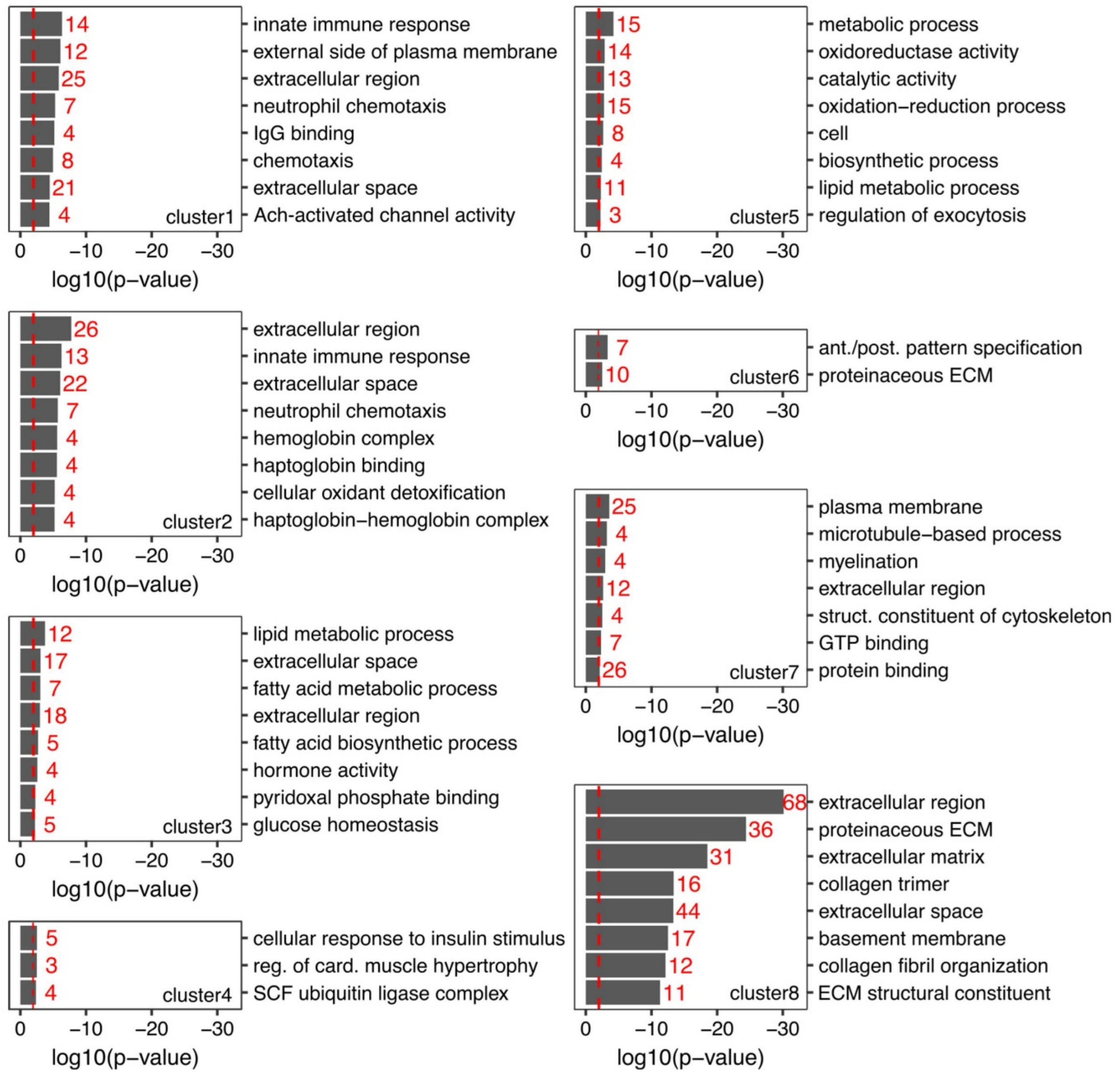


**Figure S3: Gene ontology of gene clusters.** Top-eight DAVID gene ontology terms enriched ($P < 0.01$) for genes aligned to each of the eight clusters (Fig. 3E) identified through hierarchical clustering of genes aligned with any of the four PCs described in Figure 3B. Enrichment significance threshold was set at $P < 0.01$ (gray and red dashed lines).

**Figure S4**

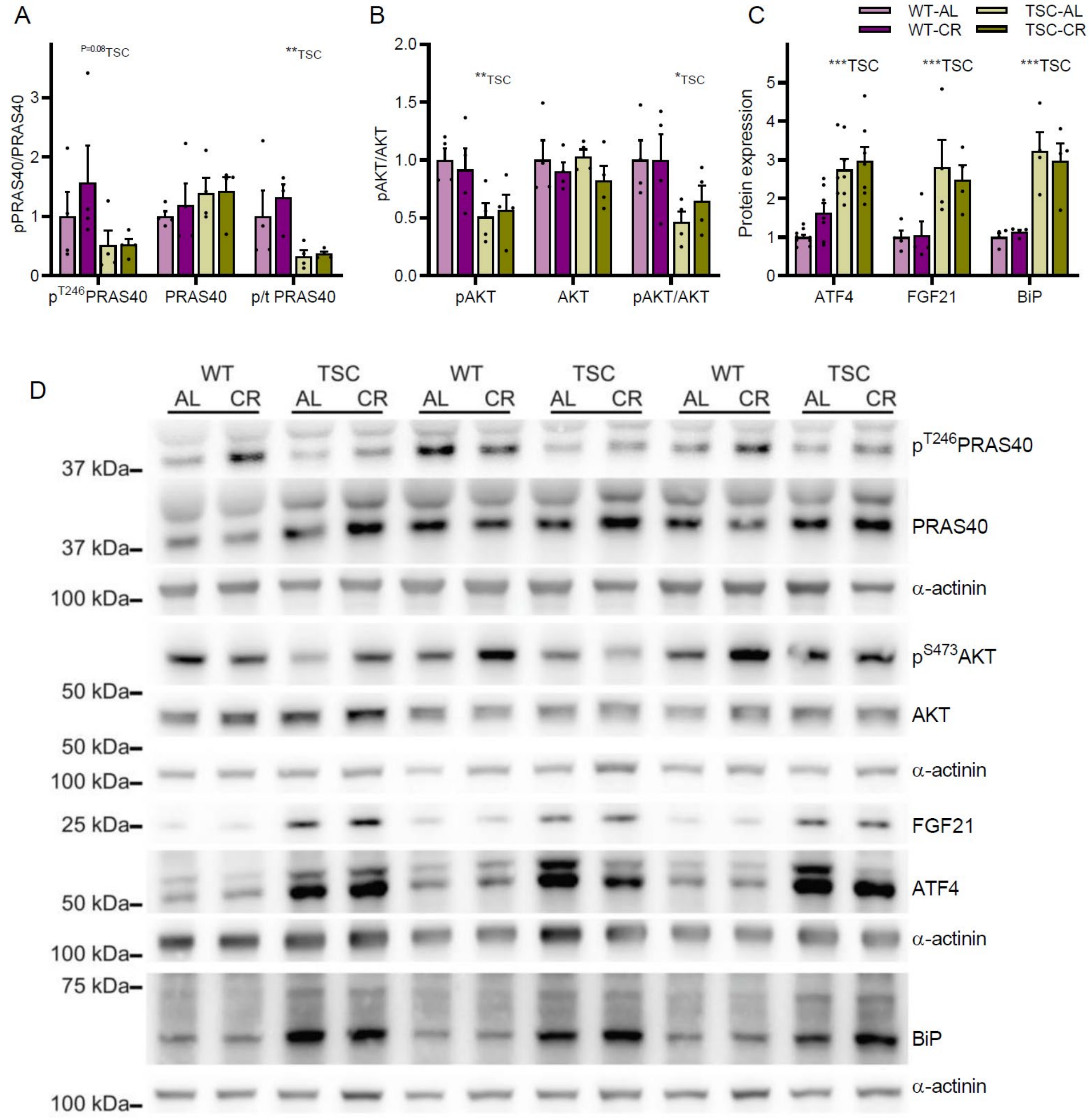


**Figure S4. CR does not alter AKT activation or ER stress induction in TSCmKO mice.** Quantification of phosphorylated and total protein levels for (A) PRAS40, (B) AKT and (C) eif2a as well as total levels of (D) ATF4, FGF21 and BiP and (E) Representative western blot images. Data are presented as mean ± SEM. Two-way ANOVAs with Tukey post hoc tests were used to compare data. *, **, and *** denote a significant difference between groups of $P < 0.05$, $P < 0.01$, and $P < 0.001$, respectively. # denotes a trend where $0.05 < P < 0.10$. Colored asterisks refer to the group of comparison.

**Figure S5**

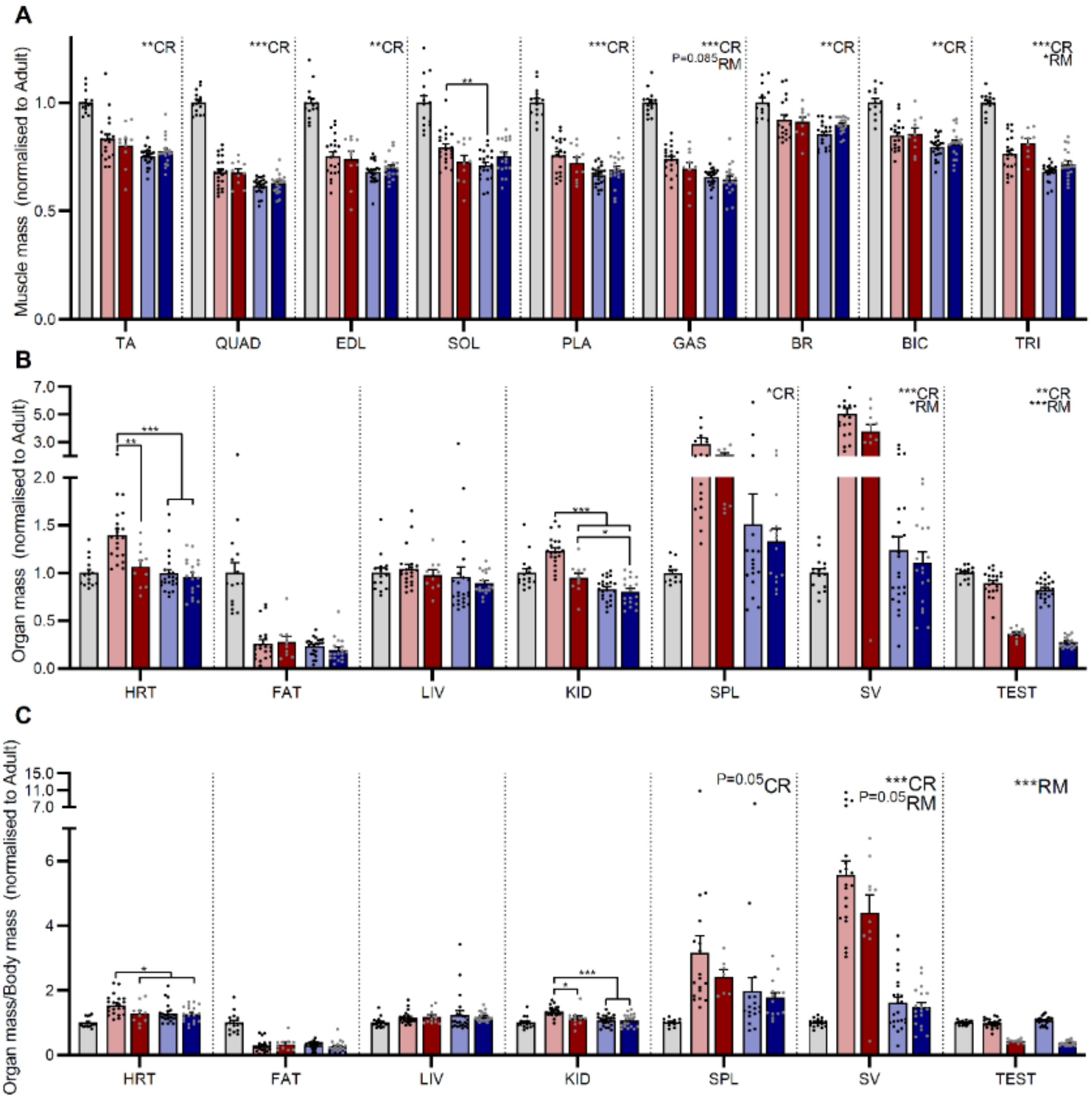


**Figure S5: CR and RM specific effects on muscle and tissue mass.** (A) Muscle mass for tibialis anterior (TA), *quadriceps* (QUAD), *extensor digitorum longus* (EDL), *soleus* (SOL), *plantaris* (PLA), *gastrocnemius* (GAS), brachioradialis (BR), biceps brachii (BIC) and *triceps brachii* (TRI) were averaged across both limbs and normalized to 10-month-old control mice. Organ mass normalized to (B) 10mCON or (C) body mass and then 10mCON, including heart (HRT), epididymal fat (FAT), liver (LIV), Kidney (KID), spleen (SPL), seminal vesicles (SV) and testicles (TEST). Data are presented as mean ± SEM. Two-way repeated- measure ANOVAs with Tukey's post hoc tests were used to compare between data. *, **, and *** denote a significant difference between groups of $P < 0.05$, $P < 0.01$, and $P < 0.001$, respectively. Colored asterisks refer to the group of comparison.
